# Synergistic activity of IL-2 mutein with tolerogenic ImmTOR nanoparticles leads to massive expansion of antigen-specific Tregs and protection against autoimmune disease

**DOI:** 10.1101/2023.05.15.540840

**Authors:** Takashi Kei Kishimoto, Max Fournier, Alicia Michaud, Gina Rizzo, Christopher Roy, Teresa Capela, Natasha Nukolova, Ning Li, Liam Doyle, Fen-ni Fu, Derek VanDyke, Peter G. Traber, Jamie B. Spangler, Sheldon S. Leung, Petr O. Ilyinskii

## Abstract

Low dose IL-2 therapy and IL-2 molecules engineered to be selective for the high affinity IL-2 receptor have been shown to expand Tregs in vivo, and, in the case of low dose IL-2 therapy, has demonstrated promising therapeutic benefit in autoimmune diseases. One of the potential limitations of IL-2 therapy is the nonselective expansion of pre-existing Treg populations rather than induction of antigen-specific Tregs, as well as potential activation of effector cells. We have recently developed biodegradable nanoparticles encapsulating rapamycin, called ImmTOR, to induce selective immune tolerance to co-administered antigens, such as immunogenic biologic drugs. Unlike Treg-selective IL-2 therapy, ImmTOR alone does not increase total Treg numbers. However, here we demonstrate that the combination of ImmTOR and an engineered Treg-selective IL-2 variant (termed IL-2 mutein) increases the number and durability of total Tregs, as well as inducing a profound synergistic increase in antigen-specific Treg when combined with a target antigen. We demonstrate that the combination of ImmTOR and an IL-2 mutein leads to durable inhibition of antibody responses to co-administered AAV gene therapy capsid, even at sub-optimal doses of ImmTOR, and provides protection in autoimmune models of type 1 diabetes and primary biliary cholangitis. ImmTOR also showed the potential to increase the therapeutic window of engineered IL-2 molecules by mitigating effector T cell expansion typically observed at higher doses of IL-2 and preventing exacerbation of disease in a model of graft-versus-host-disease. At the same time, engineered IL-2 molecules showed potential for dose-sparing of ImmTOR. Overall, these results establish that the combination of ImmTOR and an IL-2 mutein show synergistic benefit on both safety and efficacy to provide durable antigen-specific immune tolerance to mitigate drug immunogenicity and to treat autoimmune diseases.

## Introduction

The autoimmune disease field is experiencing a resurgence in the development of regulatory T cell (Treg)-focused therapies to counterbalance autoreactive effector T cells in autoimmune disease^1^. Two divergent strategies that have emerged focus on either non-selectively expanding total Tregs or inducing antigen-specific Tregs^2, 3^. Strategies to expand total Tregs frequently employ interleukin-2 (IL-2), a critical Treg growth and survival factor^4, 5^. Tregs constitutively express a high affinity trimeric IL-2 receptor, comprised of the α (CD25), β (CD122) and γ (CD132) subunits (IL-2Rαβγ, K_D_∼100 pM); whereas, effector T cells and NK cells constitutively express an intermediate affinity heterodimeric IL-2 receptor, comprised of only the β (CD122) and γ (CD132) subunits (IL-2Rβγ, K_D_∼10 nM). Low doses of IL-2 have been used clinically to selectively target the high affinity IL-2Rαβγ on Tregs^6–9^, and the difference in the affinity of IL-2 for the two receptors has also been exploited by investigators to engineer IL-2 for the treatment of either autoimmune diseases or oncology ^10, 11^. Various engineered forms of IL-2, including IL-2 muteins^12–16^, pegylated IL-2^17, 18^, IL-2-antibody complexes and fusion proteins^19–23^, and CD25-IL-2 fusion proteins ^24^, have been designed to selectively engage the high affinity IL-2 receptor, leading to biased expansion of Tregs in vivo. Large numbers of expanded and activated Tregs can provide ‘bystander suppression’ to inhibit immune responses in an antigen-independent manner, through the production of immunosuppressive cytokines, sequestration of IL-2 from effector T cells, generation of extracellular adenosine, and/or through Treg-mediated trogocytosis of co-stimulatory molecules expressed by antigen-presenting cells^25–27^. Early phase clinical trials of low dose IL-2 have demonstrated promising results in treatment of systemic lupus erythematosus (SLE), graft-versus-host disease (GVHD), and other autoimmune conditions^6–8, 28^.

Despite early successes with low dose IL-2 therapies, antigen-specific immune tolerance has been a long-standing goal for immunotherapy of autoimmune diseases, which are currently treated by systemic immunosuppressive or immunomodulatory drugs. Indeed, preclinical studies suggest that antigen-specific Tregs play a dominant role in preventing autoimmunity^29^ and are more effective than total Tregs in mitigating autoimmune disease^30^. The ratio of Treg:effector T cells is thought to be an important factor in immune homeostasis^31, 32^. Thus, the generation of large numbers of antigen-specific Tregs may be necessary to offset the large pool of pre-existing autoreactive effector T cells in the context of autoimmune disease. Autologous T cells can be engineered ex vivo to express specific T cell receptors (TCRs) in order to generate large numbers of antigen-specific Tregs; however, this strategy has the drawback of complex and costly manufacturing of personalized cell therapies. An alternative strategy is the in vivo induction of antigen-specific Tregs. Early clinical trials of antigen-specific immune tolerance strategies have shown encouraging results^33–35^.

We have developed biodegradable tolerogenic nanoparticles, termed ImmTOR, which encapsulate rapamycin, a macrolide inhibitor of the mTOR pathway^36, 37^. ImmTOR nanoparticles selectively biodistribute to the spleen and liver following intravenous injection, and they induce a tolerogenic phenotype in antigen-presenting cells that endocytose the nanoparticles^37–39^. ImmTOR has been shown to induce immune tolerance to a variety of co-administered antigens, as demonstrated by: 1) induction of antigen-specific Tregs; 2) the ability to transfer tolerance from treated mice to naïve recipients; 3) the ability to withstand subsequent challenge with the antigen alone; and 4) the ability to maintain immunological responses to unrelated antigens^36^. Therapeutic administration of ImmTOR + antigen has been shown to inhibit disease relapse in a relapsing-remitting model of experimental autoimmune encephalomyelitis (EAE)^38, 40^. In addition, ImmTOR enables repeated dosing of highly immunogenic microbial therapeutic proteins, such as a fungal-derived uricase enzyme (pegadricase), a bacterial-derived immunotoxin, and viral-derived gene therapy vectors^33, 37, 41^. ImmTOR dosed in combination with pegadricase has been shown to inhibit the formation of anti-drug antibodies in humans^33^ and is being evaluated in Phase 3 clinical trials for the treatment of chronic refractory gout (NCT04513366 and NCT04596540).

Here we describe an in vivo strategy to leverage the benefits of polyclonal Treg expansion as well as induction and expansion of antigen-specific Tregs using a combination of a Treg-selective IL-2 mutein and ImmTOR co-administered with a target antigen. This combination therapy approach produces a massive and synergistic increase in antigen-specific Tregs in vivo. When administered in combination with an adeno-associated virus (AAV) gene therapy vector, treatment with ImmTOR and an IL-2 mutein led to profound synergistic inhibition of anti-AAV antibodies, even at sub-therapeutic doses of ImmTOR. Similarly, combining ImmTOR + IL-2 mutein treatment with a nanoparticle-encapsulated autoantigen protected non-obese diabetic (NOD) mice from the development of Type 1 diabetes (T1D) and was efficacious in a mouse model of primary biliary cholangitis (PBC). In addition, ImmTOR + IL-2 mutein treatment increased the therapeutic window of engineered IL-2 by restraining effector cell activation and preventing disease exacerbation in a model of GVHD. Taken together, these results suggest that the combination of ImmTOR with a Treg-selective IL-2 molecule could be a modular and effective strategy to promote specific immune tolerance to a variety of target antigens.

## Results

### Combination of Treg-selective IL-2 + ImmTOR leads to biased Treg expansion

To investigate the effects of combining ImmTOR and a Treg-selective IL-2 on nonselective expansion of total Tregs, we employed a mouse IL-2 mutein fused to an Fc domain for extended in vivo half-life, as described by Gavin and colleagues (known as Fc.IL2m)^15, 42^. Mice treated with a single dose of Fc.IL2m alone showed a dramatic increase in total splenic Tregs, with a peak at 4 days and levels declining back to baseline by day 14 (Figure 1A, Suppl Fig 1), as previously described^13^. As expected, mice treated with ImmTOR alone showed little or no increase in total Tregs (Figure 1A, B) compared to naïve mice. However, the addition of ImmTOR amplified Treg expansion in response to Fc.IL2m. The Treg response to ImmTOR+Fc.IL2m lagged behind that observed with Fc.IL2m alone, with levels peaking approximately 7 days after treatment (Figure 1A, B). Moreover, Treg levels remained significantly increased at 14 days after treatment with ImmTOR+Fc.IL2m compared to both naïve mice and mice treated with Fc.IL2m alone. The absolute number of Tregs and the percent Tregs of total CD4+ T cells showed dose-dependent increases (up to 13-fold) following treatment with ImmTOR+Fc.IL2m compared to naïve mice (Figure 1C), and striking increases were observed in the number and percentage of Ki67^+^ proliferating Tregs and Helios^+^ stable Tregs (Figure 1C, Suppl Figure 2A). Whole animal imaging of mice expressing red fluorescence protein (RFP) under control of the Foxp3 promoter provided in situ confirmation of increased Foxp3 expression when ImmTOR was co-administered with Fc.IL2m (Figure 1D). Notably, treatment with the Treg-selective Fc.IL2m led to increases in CD8^+^ cytolytic T cells (CTL), effector CD4^+^ T cells (Teff), and natural killer (NK) cells at high doses of 18 and 27 µg (Figure 1E). In contrast, the addition of ImmTOR to Fc.IL2m suppressed the expansion of these immune effector cell subsets. The increased numbers of Tregs combined with suppression of effector cells resulted in substantially increased Treg:effector cell ratios in animals treated with ImmTOR + Fc.IL2m compared to mice treated with Fc.IL2m alone or naïve mice (Figure 1F). Similar effects of ImmTOR+Fc.IL2m treatment were observed in the liver, although the degree of Treg expansion was lower than that observed in the spleen (Suppl Figure 2B).

**Figure 1.**
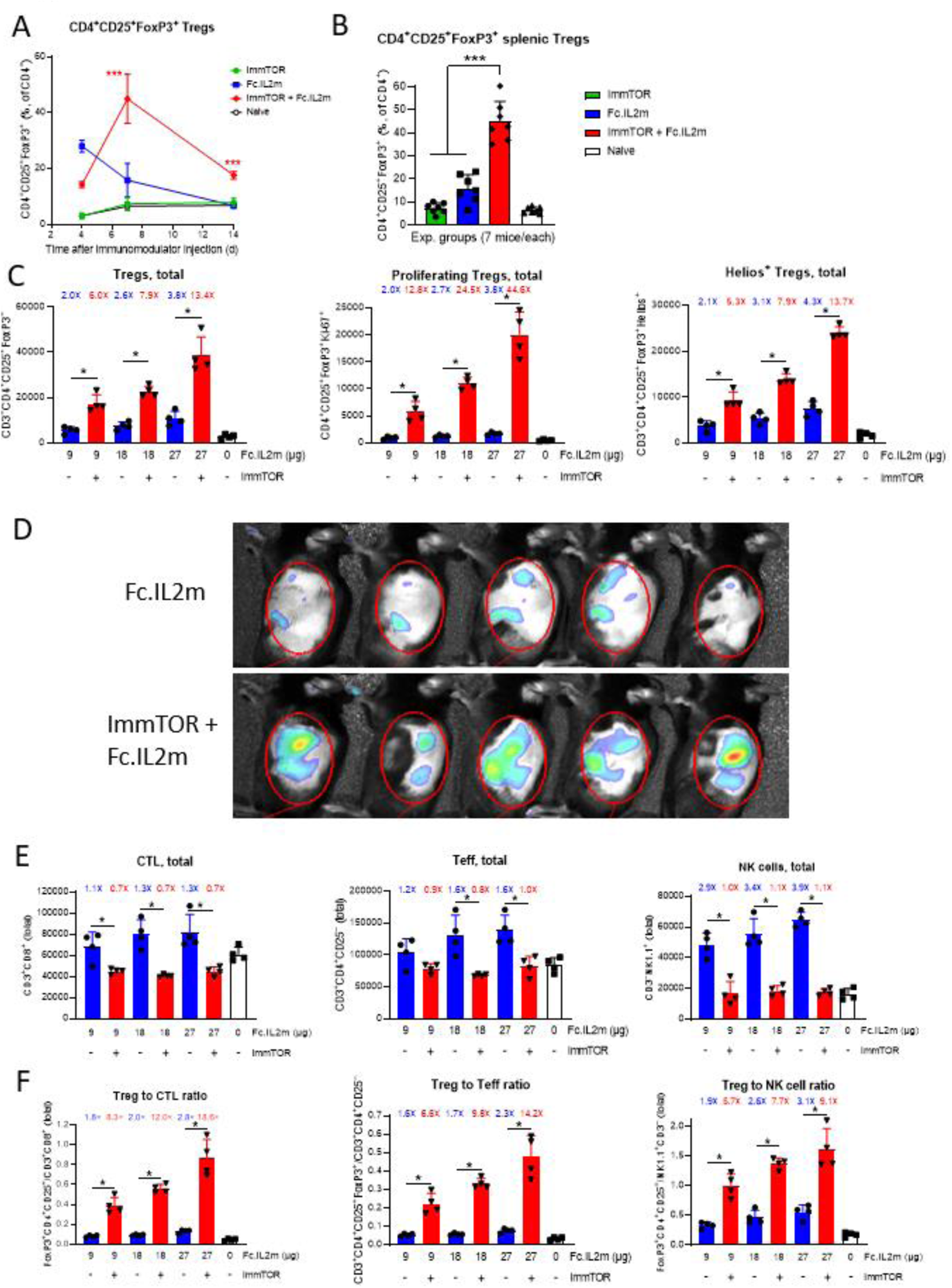
Expansion of splenic Tregs by ImmTOR and IL-2 mutein. **A**. Dynamics of Treg induction by ImmTOR, Treg-biased IL-2 mutein (Fc.IL2m) or the combination thereof. Groups of mice (n= 3-7 mice per timepoint) were treated as described, and spleens were harvested at times indicated, processed to single-cell suspension, stained, and analyzed for CD3^+^CD4^+^CD25^+^FoxP3^+^ Treg abundance by flow cytometry. This graph is a summary of four independent experiments. Error bars indicate mean +/- standard deviation (SD). **B**. Representative graph of a 7-day timepoint shown in **A**. This graph is a summary of 2 independent experiments (n=7 per group). Error bars indicate mean +/- SD. **C**. Dose-dependence of Treg induction by Fc.IL2m alone or combined with ImmTOR. Groups of mice (n=4 per cohort) were treated with ascending doses of 9, 18, or 27 µg of Fc.IL2m, alone or combined with 100 µg ImmTOR. Seven days after treatment, mice were sacrificed and harvested spleens were then evaluated for total number of Tregs, proliferating (Ki-67^+^) Tregs, and Helio^+^ stable Tregs by flow cytometry. Error bars indicate mean +/- SD. **D**. Whole animal fluorescence imaging of transgenic mice expressing mRFP under control of the Foxp3 promoter. Mice were treated with either Fc.IL2m alone (top row) or with Fc.IL2m+ImmTOR (bottom row) and analyzed 7 days after treatment. **E.** Effector cell populations induced by ascending doses of Fc.IL2m alone or in combination with ImmTOR, as described in **C**. Total numbers of CD8^+^ cytolytic T lymphocytes (CTL; CD3^+^CD8^+^), CD4^+^ T effector (Teff; CD3^+^CD4^+^CD25^−^), and NK (CD3^−^NK1.1^+^) cells are shown. Error bars indicate mean +/- SD. **F**. Ratios of total number of Tregs relative to CTL, Teff, and NK cells after treatment with ascending doses of Fc.IL2m alone or in combination with ImmTOR, as described in **C**. Ratios of the value for each experimental group vs untreated control are indicated above the bars in **C**, **E** and **F**. Representative graphs from one of two studies with similar results are shown. Error bars indicate mean +/- SD. Statistical significance: * p < 0.05, ** p < 0.01, *** p < 0.001.

We observed that circulating Fc.IL2m showed slower clearance in animals treated with ImmTOR+Fc.IL2m versus those treated with Fc.IL2m alone (Suppl Figure 3A). The clearance of Fc.IL2m correlated with the kinetics of IL-2Rα (CD25) expression, which was delayed in the ImmTOR+Fc.IL2m treated group (Suppl Figure 3B), consistent with the delayed peak of Foxp3+ Treg (Figure 1A). Circulating levels of Fc.IL2m were ∼10-fold higher at Day 4 in animals treated with ImmTOR+Fc.IL2m compared to those that received only Fc.IL2m. Treatment with ImmTOR+Fc.IL2m led to greater demethylation of the Foxp3 and EOS genes compared to treatment with Fc.IL2m alone, although the demethylation pattern of the Helios gene was similar in animals treated with ImmTOR+Fc.IL2m and those that received Fc.IL2m only (Suppl Figure 3C). ImmTOR alone showed no significant effects on methylation patterns.

As Fc.IL2m is a mouse IL-2-derived mutein, we sought to evaluate the synergy of ImmTOR with a human IL-2-derived molecule. To this end, we made use of an engineered Treg-selective human IL-2 immunocytokine, denoted F5111 IC, which is comprised of the human anti-IL-2 antibody F5111^20^ covalently tethered to human IL-2^22^. F5111 IC has been shown to potently and selectively stimulate the high affinity IL-2R resulting in robust in vitro activation and in vivo expansion of Tregs^22^. We evaluated the activity of F5111 IC and ImmTOR in immunodeficient NOD SCID gamma mice 2-3 weeks after engraftment with human peripheral blood mononuclear cells (HuPBMC). The HuPBMC mice, which are prone to develop GVHD, showed marked expansion of Treg, CD8^+^ T cells, and NK cells after F5111 IC treatment (Figure 2A). Although the mice did not show signs of GVHD at the time of treatment, the expansion of effector cells may reflect sub-clinical inflammation of HuPBMC in response to host mouse antigens. Importantly, the addition of ImmTOR to F5111 IC treatment enabled expansion of Treg but inhibited the expansion of CD8^+^ T cells and NK cells (Figure 2A). We next evaluated the effects of F5111 IC and ImmTOR in a model of GVHD in which host mice were irradiated prior to transfer of HuPBMC, which accelerates disease course. F5111 IC alone exacerbated disease, leading to increased mortality, while the combination of ImmTOR + F5111 IC prolonged survival and improved disease scores (Figure 2B and Suppl Figure 4). Treatment with ImmTOR alone was similarly efficacious, but the combination with F5111 IC showed a trend to better durability of response.

**Figure 2.**
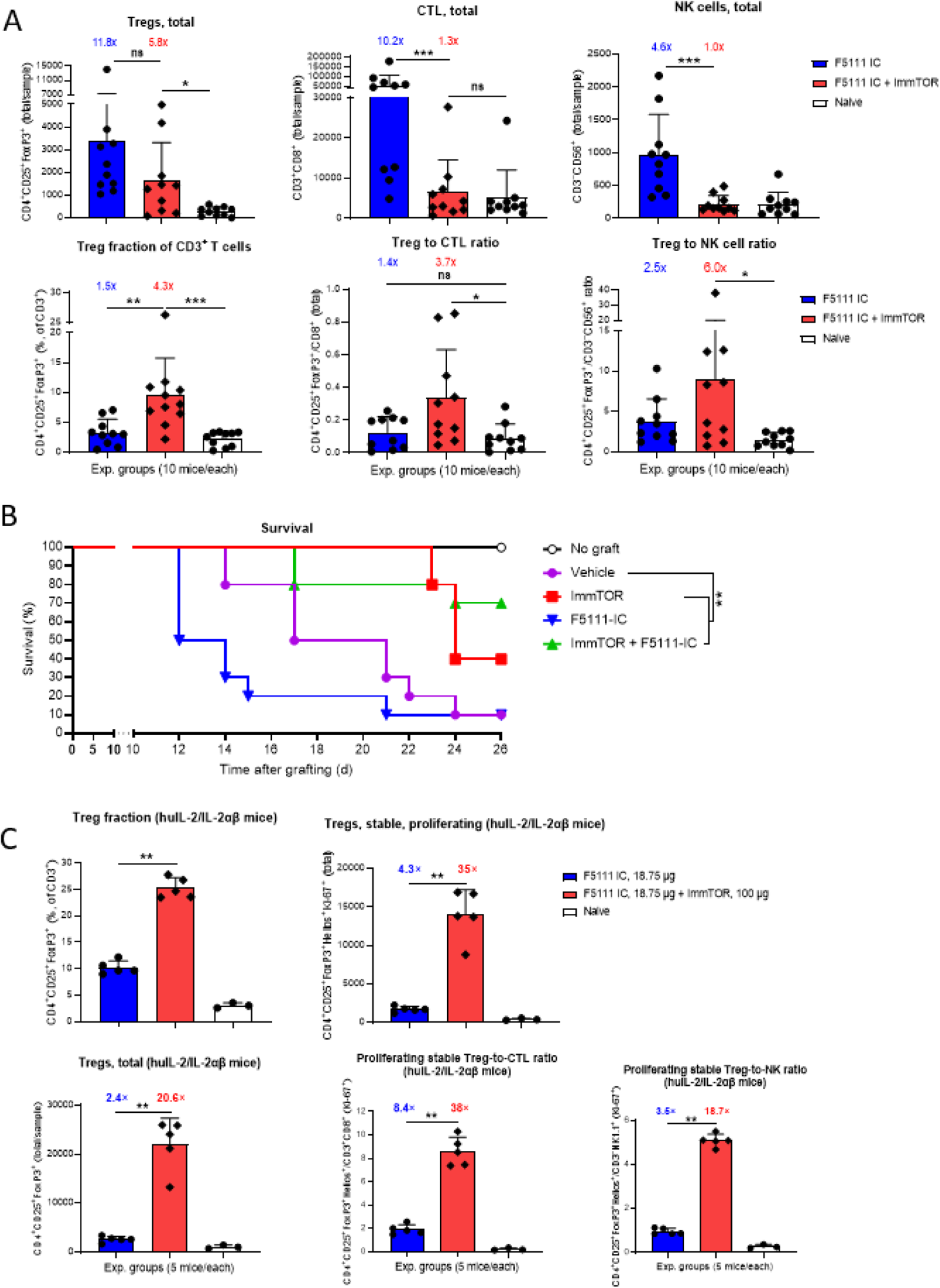
Induction of Tregs by ImmTOR and human IL-2/anti-IL-2 antibody fusion protein in humanized mice. Mice were treated with F5111 IC (18.75 µg) alone or combined with ImmTOR (100 µg), and splenocytes were harvested at 7 days post treatment and analyzed by flow cytometry. **A**. Human PBMC-engrafted NSG (huPBMC) mice were treated at 1.5-3 weeks after PBMC engraftment. Treg (CD3^+^CD4^+^CD25^+^FoxP3^+^), CTL (CD3^+^CD8^+^), and NK cell (CD3^−^CD56^+^) populations are presented as absolute cell numbers, percent Tregs out of total T cells, and relative ratios of Treg:effector cells. A summary of 3 experiments using different PBMC donors is shown (n=10 per group). Error bars indicate mean +/- SD. **B**. ImmTOR mitigates disease exacerbation by F5111 IC and prolongs survival in a HuPBMC model of GVHD. NSG mice were irradiated and then reconstituted with 1×10^7^ human PBMC. The next day, mice were treated with a single dose of saline, ImmTOR (100 µg), F5111 IC (9 µg), or the combination. Control animals were irradiated but did not receive HuPBMC. **C.** Transgenic mice expressing human IL-2, IL-2Rα and IL-2Rβ mice were treated as described (5 animals/group) or left untreated (3 animals/group). Treg, Helios^+^ stable Treg, CTL, and NK total and proliferating cell populations are shown as fractions, absolute cell numbers, or relative ratios. A representative experiment of 2 independent studies that resulted in a similar outcome is shown. Error bars indicate mean +/- SD. Ratios of the value for each experimental group vs untreated control are indicated above the bars in **A** and **C**. Statistical significance: * p < 0.05, ** p < 0.01, *** p < 0.001.

We next assessed the effects of F5111 IC in a non-disease setting using engineered knock-in mice expressing human IL-2Rαβ, which can form functional IL-2 receptors with endogenously expressed mouse IL-2Rγ. F5111 IC induced robust Treg expansion in the engineered human IL-2Rαβ mice without substantial expansion of CD8^+^ T cells (Figure 2C). The addition of ImmTOR to F5111 IC resulted in a corresponding synergistic expansion of Tregs, similar to that observed in wild-type mice treated with ImmTOR+Fc.IL2m. Collectively, the results of our mouse and human Treg expansion studies demonstrate that combination treatment with ImmTOR and Treg-selective IL-2 molecules induces selective promotion of Treg proliferation, synergistically enhancing the activity of the Treg-selective IL-2 molecules alone while restraining effector cell activation.

### ImmTOR+Fc.IL2m treatment ameliorates autoimmune hepatitis

The activity of the ImmTOR+Fc.IL2m was evaluated in a model of autoimmune hepatitis induced by systemic administration of the concanavalin A, a lectin that causes polyclonal lymphocyte activation and hepatic infiltration of activated immune cells. Previous studies have shown that Treg depletion with anti-IL-2Rα antibodies exacerbated disease while adoptive transfer of Treg ameliorated disease^43^. Both ImmTOR and Fc.IL2m monotherapies inhibited infiltration of activated effector T cells, and the combination treatment led to a further reduction in T cell infiltrates (Figure 3A). Similar, though more modest, reductions were observed in activated NK cells; whereas reductions in activated NKT cells, neutrophils, and macrophages were primarily mediated by ImmTOR (Suppl Figure 5). Both ImmTOR and Fc.IL2m reduced production of serum interferon-γ (IFN-γ) and, to a lesser extent, of CXCL1 chemokine (Figure 3B). Combination treatment further reduced the levels of both IFN-γ and CXCL1, whereas reductions in IL-6 were primarily mediated by ImmTOR. ImmTOR+Fc.IL2m administration also induced increased production of FGF21, a hepatoprotective stress-response growth factor (Figure 3C). Overall, this model illustrates the therapeutic potential for ImmTOR+Fc.IL2m combination therapy in autoimmune hepatitis.

**Figure 3.**
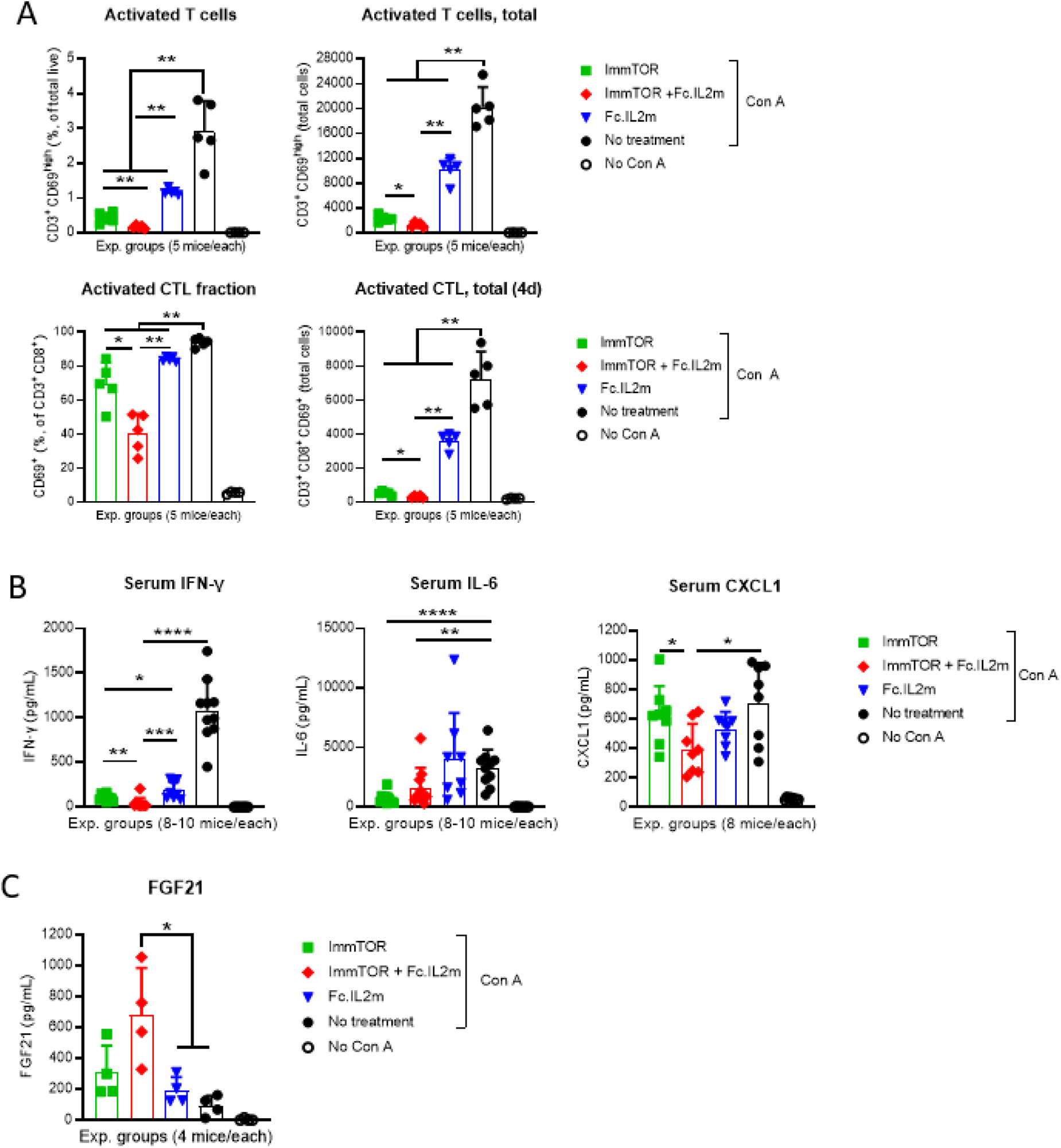
Efficacy of ImmTOR + Treg-biased IL-2 mutein in a concanavalin A-induced model of autoimmune hepatitis. **A.** Female C57BL/6 mice (n=5 per cohort) were either left untreated or treated with ImmTOR (200 µg) and Fc.IL2m (9 µg) individually or in combination. Mice were challenged 4 days later with 12 mg/kg of concanavalin A (Con A) or left unchallenged (no Con A). At 12 hours after Con A challenge, serum was drawn for cytokine analysis and livers were harvested and hepatic T cells were assessed by flow cytometry. **A**. Activated (CD69^+^) and highly activated (CD69^high^) T cells (CD3^+^) and CTL (CD3^+^CD8^+^) are shown either as fractions of total or as absolute cell numbers. A representative experiment of 4 studies that resulted in similar outcomes is shown. Error bars indicate mean +/- SD. **B**. Mice were treated as in **A**. Serum IFN-γ, IL-6, and KC/GRO levels at 12 hours after Con A challenge (summary of 2 independent experiments, n=8-10 per group). Error bars indicate mean +/- SD. **C**. FGF21 serum levels prior to and after Con A challenge. A representative experiment of 4 studies that resulted in similar outcomes is shown (n=4 per group). Error bars indicate mean +/- SD. Statistical significance: * p < 0.05, ** p < 0.01, *** p < 0.001, **** p < 0.0001.

### ImmTOR+Fc.IL2m leads to synergistic induction and expansion of antigen-specific Treg

ImmTOR has been shown to induce antigen-specific Treg to co-administered antigens^36, 37^. We therefore sought to test whether ImmTOR+Fc.IL2m could enhance the induction of antigen-specific Treg when co-administered with antigen. In these experiments, we utilized ovalbumin (OVA) as the target antigen following adoptive transfer of OVA-specific OT-II CD4^+^ T cells. As expected, ImmTOR+OVA did not expand pre-existing host Treg but induced OT-II Foxp3^+^ Treg (Figure 4A). In contrast, Fc.IL2m+OVA expanded host Tregs but did not have a significant effect on OT-II Tregs. The triple combination of ImmTOR+Fc.IL2m+OVA led to a profound synergistic expansion of OVA-specific OT-II Tregs, while also increasing total host Treg compared to Fc.IL2m+OVA or ImmTOR+OVA treatment alone. ImmTOR+Fc.IL2m without OVA had only a modest effect on OT-II Tregs, consistent with previous findings that the target antigen must be co-administered with ImmTOR to induce antigen-specific Tregs^37^. Increasing amounts of OVA led to a dose-dependent increase in OT-II Treg in response to ImmTOR+Fc.IL2m, plateauing at approximately 100 µg OVA (Figure 4B). As anticipated, there was no significant effect of increasing amounts of OVA on the host Treg response.

**Figure 4.**
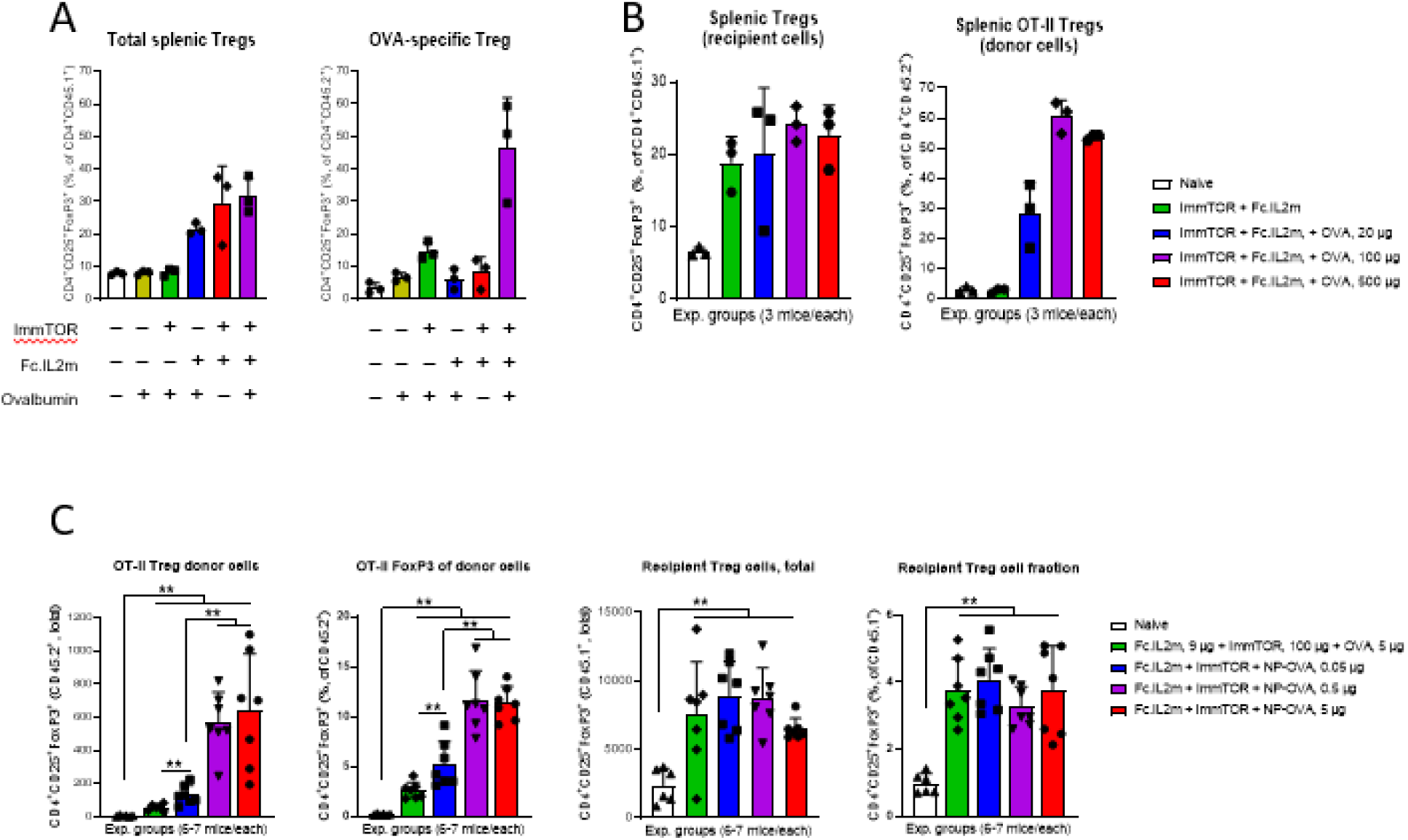
Induction of OVA-specific Tregs by combination of ImmTOR, IL-2 mutein, and ovalbumin. OVA-specific OT-II T cells from CD45.2^+^ donors were adoptively transferred into CD45.1^+^ C57BL/6 recipient mice (n=3 per group) and 24 hours later treated with different combinations of ImmTOR (100 µg), Fc.IL2m (9 µg), and either free ovalbumin (OVA, 5-500 µg) or nanoparticle-encapsulated OVA (NP-OVA, 0.05-5 µg) or left untreated. At 7 days after treatment, spleens were harvested and analyzed by flow cytometry. **A**. Total splenic and OVA-specific Tregs induced by combinations of ImmTOR, Fc.IL2m, and free OVA (n=3/group). Error bars indicate mean +/- SD. **B**. Antigen dose-dependent induction of OVA-specific donor Tregs by the combination of ImmTOR, Fc.IL2m, and free OVA. The concurrent expansion of recipient Tregs was unaffected by OVA. Total numbers of CD45.1^+^ (recipient) and CD45.2^+^ (OT-II donor) Tregs are shown (n=3 per group). Error bars indicate mean +/- SD. **C.** Antigen dose-dependent induction of OVA-specific donor or recipient Tregs by the combination of ImmTOR, Fc.IL2m, and either free OVA or NP-OVA. Total number of CD45.1^+^ (recipient) and CD45.2^+^ (OT-II donor) Tregs and the fractions of total cells are shown (n=6-7 per group). Error bars indicate mean +/- SD. Statistical significance: ** p < 0.01.

Free OVA is expected to biodistribute widely, whereas ImmTOR has been shown to biodistribute selectively to the spleen and liver^37–39^. Thus, only a small proportion of the total free OVA dose is expected to co-localize with the splenic and hepatic antigen-presenting cells that endocytose ImmTOR. We therefore examined whether formulating ovalbumin in nanoparticles would improve the efficiency of OT-II Treg expansion by increasing co-localization with ImmTOR. Nanoparticles encapsulating 0.05 µg OVA (NP-OVA) enabled significantly more OT-II Treg induction and expansion in response to ImmTOR+Fc.IL2m treatment than did free OVA at 100-fold higher doses (Figure 4C). Maximal Treg expansion was observed at 0.5 µg nanoencapsulated OVA, with further increase of NP-OVA providing no additional Treg elevation. Taken together, these OT-II mouse studies established that ImmTOR+Fc.IL2m treatment leads to robust antigen-specific Treg expansion when co-administered with the target antigen, particularly with nanoencapsulated antigen.

### ImmTOR+Fc.IL2m treatment synergistically inhibits anti-AAV antibody responses

Combination ImmTOR+Fc.IL2m treatment was assessed for the ability to inhibit antibody responses to an adeno-associated virus (AAV) gene therapy vector. Mice immunized with two doses of AAV8 on Day 0 and Day 56 developed rapid and robust anti-AAV immunoglobulin G (IgG) antibody responses (Figure 5). As previously reported^44, 45^, ImmTOR was shown to inhibit anti-AAV antibody responses when co-administered with AAV vectors at a dose of 200 µg; however, some animals exhibited late breakthrough of anti-AAV antibodies at Day 91 (Figure 5). Suboptimal doses of 50 or 100 µg ImmTOR resulted in earlier breakthrough of antibody responses. Similarly, Fc.IL2m co-administered with AAV on Days 0 and 56 showed weak modulation of AAV immunogenicity, with antibodies detected as early as 12 days after immunization. In contrast, ImmTOR+Fc.IL2m combination treatment resulted in complete inhibition of anti-AAV antibodies, even at sub-optimal doses of ImmTOR. Anti-AAV IgG levels were significantly lower in animals treated with ImmTOR+Fc.IL2m compared to those treated with ImmTOR alone from day 33 onwards (Suppl Table ST1). These results indicate that combining ImmTOR with Fc.IL2m provides synergistic benefit for the efficacy and durability of AAV immunogenicity mitigation.

**Figure 5.**
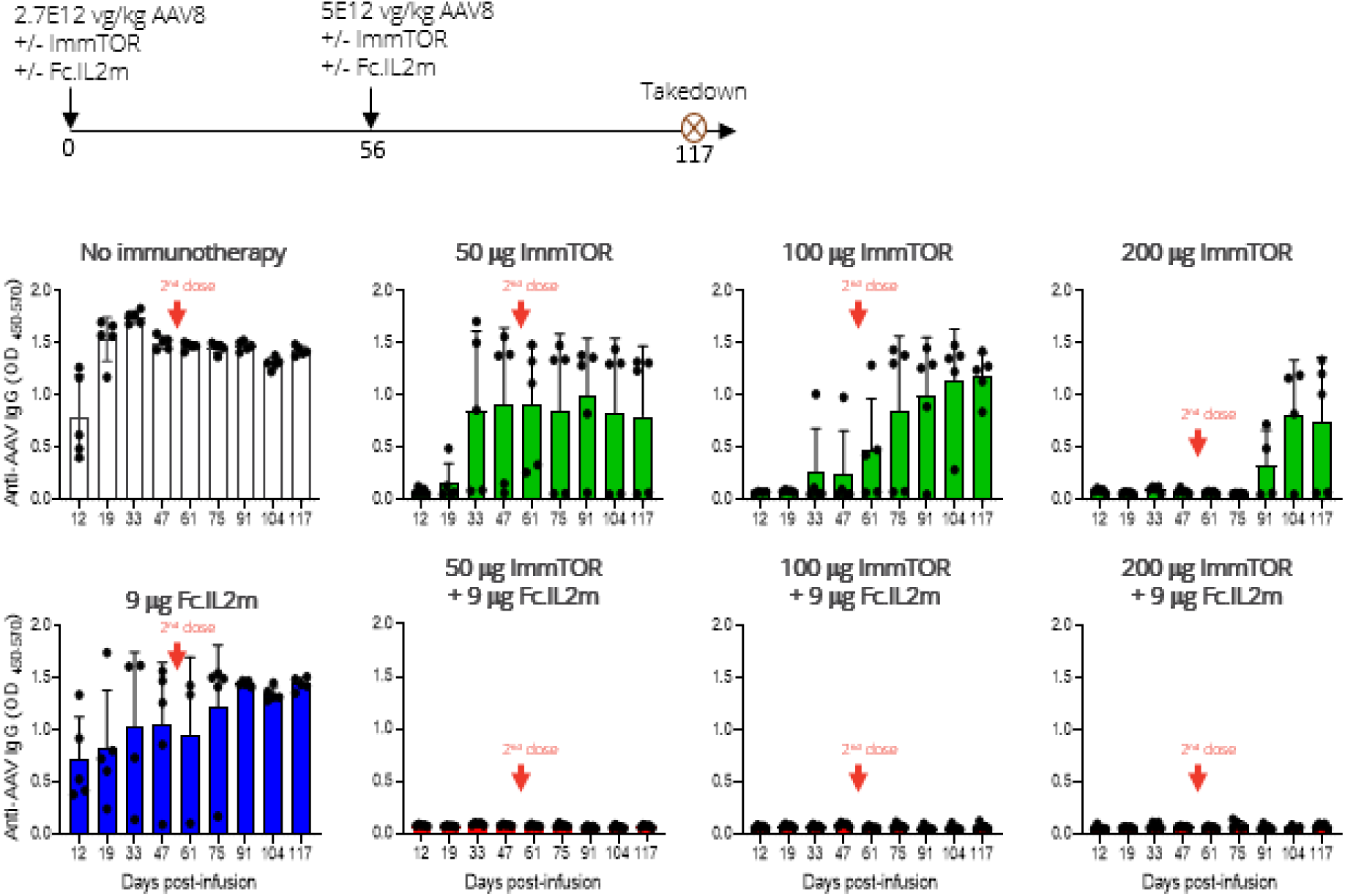
Mitigation of antibody response to AAV vector by combination treatment with ImmTOR and IL-2 mutein. C57BL/6 mice (5 per group) were injected with AAV8 on Day 0 (2.7E12 vg/kg) and Day 56 (5.0E12 vg/kg) either alone or in combination with ImmTOR (50-200 µg) and/or Fc.IL2m (9 µg). Animals were bled at the indicated timepoints indicated, and serum was analyzed for the presence of anti-AAV8 IgG antibody by ELISA. Timing of the second AAV8 administration is shown by arrows. Error bars indicate mean +/- SD. Statistical analyses are shown in Supplementary Table ST1.

Similar synergistic effects of ImmTOR+Fc.IL2m were observed using a high vector dose of 5E13 vg/kg AAV8, showing control of anti-AAV IgG development for more than 4 months after AAV inoculation (Suppl Fig 6A). Immune phenotyping of splenocytes also reflected the synergistic effect of ImmTOR and Fc.IL2m. There were trends towards increased numbers of total splenic CD8^+^ T cells, CD4^+^ effector T cells, and B cell plasmablasts four days after administration of a high vector dose of AAV alone (Suppl Fig 6B). Robust expansion of total Tregs was observed at Day 4 when Fc.IL2m was co-administered with AAV; however, this was also accompanied by substantial expansion of CD8^+^ T cells and plasmablasts. The combination of ImmTOR with Fc.IL2m co-administered with high dose AAV induced robust expansion of total Tregs at Days 4 and 7 while inhibiting effector T cell expansion, resulting in significantly elevated Treg:CTL ratios (Suppl Fig 6B, Suppl. Table ST2).

### Efficacy of ImmTOR+Fc.IL2m in autoimmune disease is enhanced by co-administration of nanoencapsulated antigen

We next evaluated the ability of ImmTOR + Fc.IL2m to prevent type 1 diabetes in the NOD mouse model. NOD mice were administered 4 monthly treatments starting at 8 weeks of age (Figure 6A). ImmTOR and ImmTOR+Fc.IL2m were administered in the absence or presence of nanoparticle-entrapped hybrid diabetes peptide 6.9^46^ (NP-HIP6.9). Whereas 7 out of 10 mice in the control group progressed to diabetes by week 30, all treated groups showed evidence of disease protection (Figure 6A and 6B). The cohort treated with ImmTOR+Fc.IL2m combined with NP-HIP6.9 was the only group that did not have a single diabetic mouse by week 33. Notably, even in the absence of co-delivered antigen, the ImmTOR+Fc.IL2m combination protected 9 out of 10 mice from diabetes at week 33.

**Figure 6.**
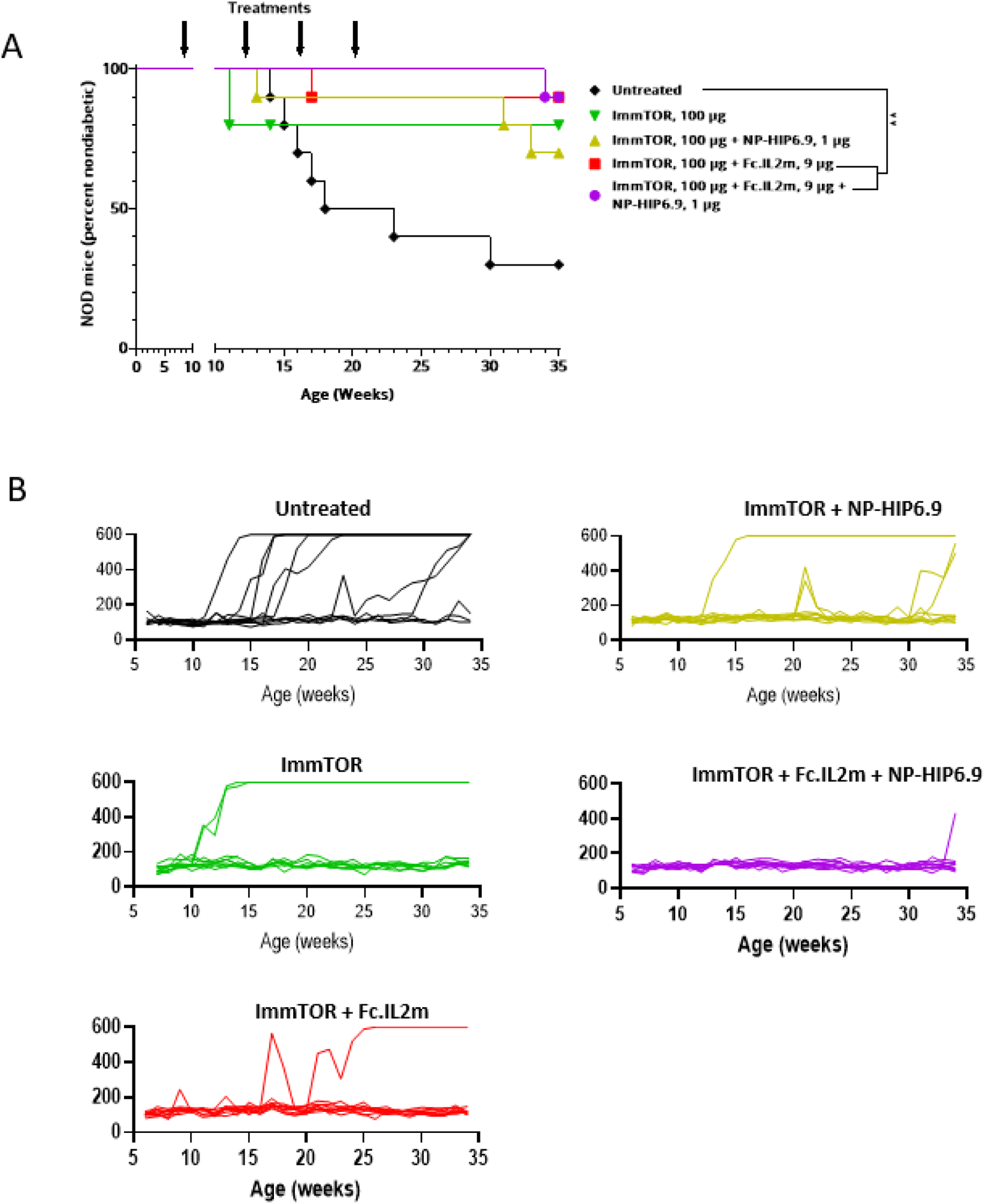
Diabetes prevention by combination treatment with ImmTOR, IL-2 mutein, and nanoparticle-encapsulated hybrid insulin peptide 6.9 (NP-HIP6.9). Female NOD mice (n=10 per group) were left untreated or treated with ImmTOR (100 µg) only or ImmTOR and Fc.IL2m (9 µg), in the absence or presence of NP-encapsulated hybrid insulin peptide 6.9 (NP-HIP6.9, 1 µg) starting at week 8 of age (4 treatments at 4-week intervals, shown as arrows). Mice were monitored up to 35 weeks of age. Blood glucose was measured weekly, and mice scoring >250 mg/dL on 2/3 successive measurements were considered diabetic and those scoring >500 mg/dL twice or >600 mg/dL once were terminated. Fractions of diabetic mice (**A**) and individual mouse blood glucose levels (**B**) are shown with statistical significance indicated. Statistical significance: ** p < 0.01.

We also assessed the activity of ImmTOR and Fc.IL2m in NOD.C3C4 mice, which spontaneously develop an autoimmune disease of the liver which closely resembles primary biliary cholangitis (PBC)^47, 48^. The primary T cell epitope has been mapped to a peptide in the inner lipoyl domain of the E2 component of the pyruvate dehydrogenase complexes (PDC-E2-ILD)^47^. Mice were treated with three monthly doses of ImmTOR, ImmTOR+Fc.IL2m or ImmTOR+Fc.IL2m combined with nanoencapsulated PDC-E2-ILD antigen (NP-PDC-E2-ILD) (Figure 7A). Treatment with ImmTOR+Fc.IL2m significantly reduced bile duct epithelial degeneration, biliary hyperplasia and liver inflammation (Figure 7B). Co-administration of NP-PDC-E2-ILD provided additional benefit. Liver histology showed striking biliary pathology, with marked peri-biliary mononuclear cell infiltrates, biliary hypercellularity and ductular ectasia in both female (Figure 7C-F) and male mice (Figure7 G-J). Treatment with ImmTOR (Figure 7D and 7H), ImmTOR+Fc.IL2m (Figure 7E and 7I), and ImmTOR+Fc.IL2m+NP-PDC-E2-ILD (Figure 7F and 7J), showed progressive improvement of all histologic features, with the triple therapy showing only minimal residual disease pathology. Collectively, these AAV immunogenicity and T1D and PBC autoimmune disease models highlight the therapeutic promise of combining ImmTOR with a Treg-selective IL-2 molecule, particularly in the context of antigen co-administration.

**Figure 7.**
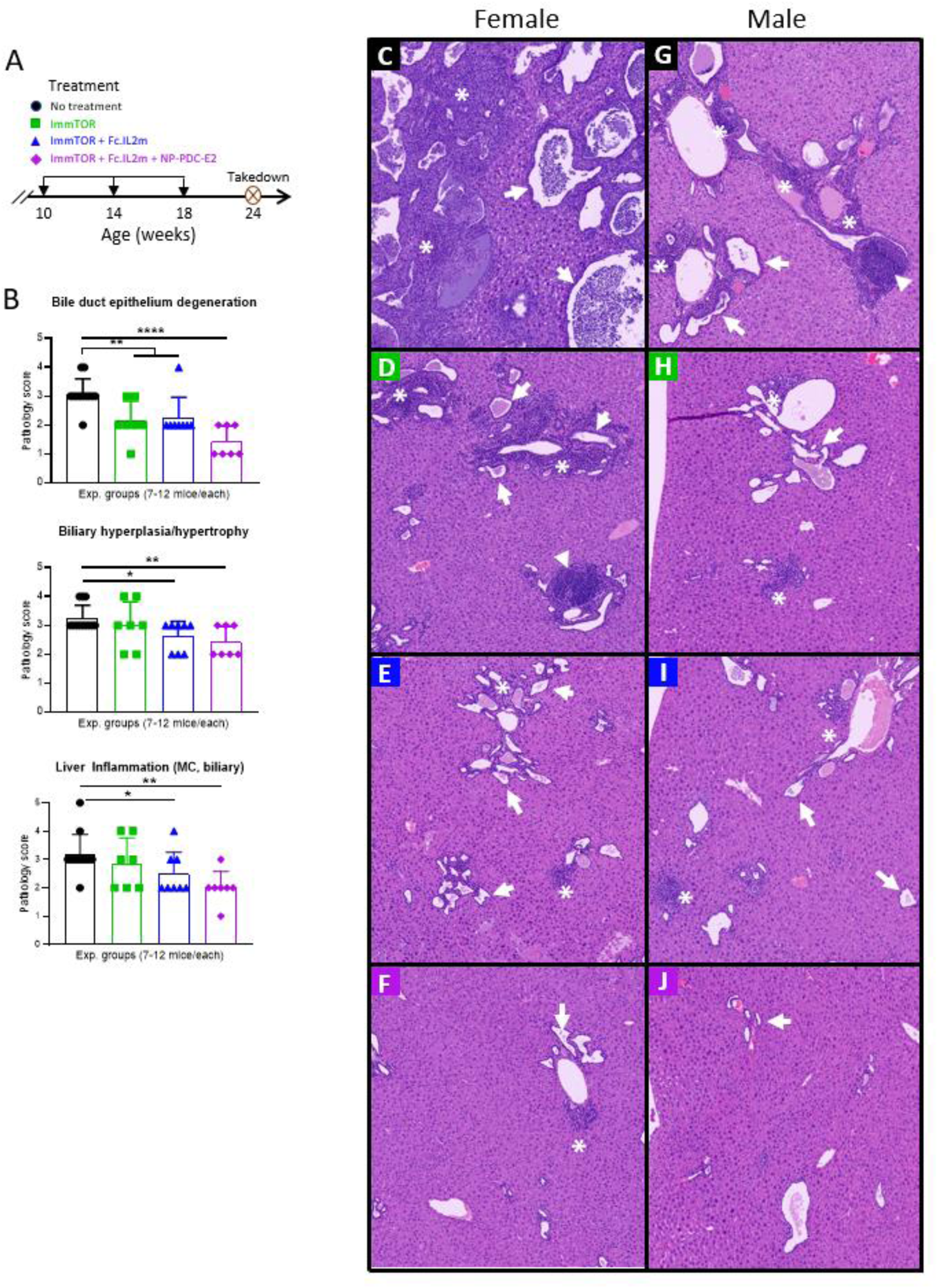
Efficacy of ImmTOR + IL-2 mutein with and without nanoparticle-encapsulated PDC-E2 antigen in a mouse model of primary biliary cholangitis. **A.** Schedule of treatment of NOD.c3c4 mice. Mice were treated with three monthly doses as indicated starting at 10 weeks of age. **B.** Disease scores. Paraffin-embedded liver sections were stained with hematoxylin & eosin of NOD.c3c4 mice and analyzed by an independent veterinary pathologist. Bile duct degeneration, biliary hypertrophy, and liver inflammation were graded on a 5-point severity scale as described in Materials and Methods. **C-J.** Histology. Representative histology sections from female **(C-F)** and male **(G-J)** mice are shown at 5x magnification. **C.** Untreated female. Vast majority of bile ducts show marked ectasia, some with intraluminal accumulations of neutrophils (arrow), and dense biliary and intrahepatic mononuclear inflammatory infiltrate (*). **D.** ImmTOR-treated female. Marked biliary mononuclear cell inflammation (*) occasionally forming follicles (arrowhead) and multifocal duct ectasia (arrow). **E.** ImmTOR-IL-treated female. Biliary mononuclear cell inflammation with small foci of peri-biliary hypercellularity (*) and multifocal duct ectasia (arrow). **F.** ImmTOR-IL plus NP/PDC-E2.ILD-treated female. Mild biliary mononuclear cell inflammation with small foci of peri-biliary hypercellularity (*) and mild biliary hyperplasia or bile duct ectasia (arrow). **G.** Untreated male. Marked biliary mononuclear cell inflammation with densely cellular accumulations surrounding bile ducts (*), with follicle formation (arrowhead), moderate bile duct hyperplasia and bile duct ectasia (arrow). **H.** ImmTOR-treated male. Biliary mononuclear cell inflammation with few foci of peri-biliary hypercellularity (*) and multifocal duct ectasia (arrow). **I.** ImmTOR-IL-treated male. Biliary mononuclear cell inflammation with small foci of peri-biliary hypercellularity (*) and multifocal duct ectasia (arrow). **J.** ImmTOR-IL plus NP/PDC-E2.ILD-treated male. Minimal biliary mononuclear cell inflammation, rare foci of peri-biliary hypercellularity and, minimal biliary hyperplasia or bile duct ectasia (arrow).

## Discussion

We describe here profound synergistic activity between ImmTOR nanoparticles carrying rapamycin and engineered IL-2 molecules that selectively activate Tregs to expand the number and durability of total Treg population, as well as in inducing and expanding antigen-specific Tregs in the presence of a co-administered target antigen. This combination therapy leverages the large body of preclinical and clinical work showing that Treg-directed IL-2 therapy can be used to selectively expand pre-existing Tregs in vivo, particularly memory and activated Tregs, for the treatment of a wide range of autoimmune diseases,^4, 5, 49^ while adding the ability to induce autoantigen-specific Tregs, which have been shown to be more effective than Tregs that are not disease-specific in animal models of autoimmune disorders^30^. While expansion of total Tregs has shown benefit in the treatment of autoimmune diseases in animals and in humans^6–9^, the ability to induce Tregs of novel antigen specificity may have additional benefits in autoimmune diseases driven by antigens for which natural thymic-derived Tregs are unlikely to exist. These include neoantigens created by post-translational modification of self-antigens, such as hybrid antigens in the case of type 1 diabetes and citrullinated antigens in the case of rheumatoid arthritis, as well as diseases driven by environmental antigens, such as transglutaminase-modified gluten proteins in the case of celiac disease^32, 46, 50–54^. In addition, our technology offers the possibility of inducing of antigen-specific immune tolerance to foreign therapeutic proteins, such as microbial enzymes or viral gene therapy vectors, which would be desirable to mitigate immunogenicity that can compromise the safety and/or efficacy of these treatments^55–57^.

Tregs are highly responsive to in vivo therapy with low dose IL-2 or engineered Treg-selective IL-2. Curiously, purified Treg do not proliferate in response to IL-2 in vitro^58^, suggesting that other signals, such as endogenous antigens presented by antigen-presenting cells and/or co-stimulation, may be required for Tregs to respond to IL-2. Although Tregs are primed to respond to IL-2 in vivo, they do not respond to in vivo treatment with rapamycin-loaded nanoparticles (ImmTOR) alone (Figure 4A). Rapamycin has been used to expand Tregs ex vivo, but the culture conditions require either the addition of exogenous IL-2 or polyclonal activation of total T cells, which are a potential source of IL-2^59, 60^. We have demonstrated that in vivo administration of ImmTOR with a target antigen promotes induction of antigen-specific Treg but does not impact total Treg abundance^37, 38, 40^ (Figure 4A). The number of antigen-specific Tregs induced by ImmTOR is limited, perhaps because production of endogenous IL-2 is limited to antigen-specific effector cells responding to the same antigen. We hypothesized that the number and durability of antigen-specific Tregs could be further enhanced by the addition of exogenous Treg-selective IL-2. Our in vivo results are consistent with in vitro studies showing that the addition of rapamycin increased Treg abundance in T cells treated with IL-2 or activated with anti-CD3 and anti-CD28 antibodies^59^. Mechanistic studies indicate that the addition of rapamycin to IL-2 stimulated T cells is due to increased frequency of Foxp3 expression rather than selective proliferation of Foxp3^+^ cells^59^.

Interestingly, a single administration of Fc.IL2m with ovalbumin showed little or no specific expansion of adoptively transferred OVA-specific T cells (Figure 4A). Recently, Gavin and colleagues demonstrated that multiple cycles of treatment with an Fc.IL2m combined with OVA in the form of a dendritic cell-targeted anti-DEC205-OVA fusion protein were required to induce and expand OVA-specific OT-II Tregs^42^. Each cycle of treatment consisted of initially expanding total Tregs, including adoptively transferred OT-II cells, using an IL-2 mutein alone, followed 2-4 days later with anti-DEC205-OVA. The rationale for this approach was to provide a selective survival strategy for the OT-II Tregs by activating them with antigen at the peak of total Treg expansion. A single cycle of IL-2 mutein followed by DEC205-OVA treatment showed no significant increase in OT-II Tregs 6 days after initiation of treatment; however, three weekly cycles of treatment increased OT-II Tregs from a baseline of ∼ 3% to 18%. These results are consistent with our results showing that the same engineered IL-2 mutein showed little or no expansion of antigen-specific OT-II T cells after a single dose. In contrast, a single treatment of ImmTOR + IL-2 mutein + OVA increased OT-II Tregs from a baseline of ∼3% to ∼45% by 7 days after treatment (Figure 4A). Daniel et al. showed that everolimus, a second generation rapalogue could also expand adoptively transferred antigen-specific Tregs when combined with the high affinity IL-2R-biased anti-IL-2 antibody JES6-1 complexed with IL-2 + antigen^61^. In this study, daily dosing of everolimus at 100µg/day for 14 days, a regimen that is typically used to mediate chronic immune suppression to prevent graft rejection, was required for Treg expansion. In contrast, our results show that a single dose of 100 µg ImmTOR was sufficient to induce antigen-specific Treg expansion when combined with an IL-2 mutein. Our results therefore suggest that combination treatment with ImmTOR and Treg-biased IL-2 molecules may allow for more efficient expansion of antigen-specific Treg compared to previous strategies.

Combination therapies are warranted for complex and serious diseases, but these approaches are often limited by additive or synergistic toxicity. ImmTOR co-administration with Treg-selective IL-2 may represent a rare exception in which combination therapy is less toxic than the individual components. The dose limiting toxicity of ImmTOR in human clinical trials has been stomatitis, a common rapamycin-associated side-effect^33^. Evaluation of the combination of ImmTOR with an IL-2 mutein showed synergistic activity in preventing antibody responses to an AAV gene therapy vector, even at subtherapeutic doses of ImmTOR (Figure 5), suggesting that the addition of an IL-2 mutein could allow for dose-sparing of ImmTOR. Conversely, the primary concern of IL-2-based therapies is the activation and expansion of effector cells, including CD4^+^ and CD8^+^ effector T cells, as well as NK cells^62^. Rapamycin, the active component of ImmTOR, is known to inhibit effector cell proliferation, while being permissive for Treg proliferation, and we observed that the addition of ImmTOR to high-dose IL-2 mutein therapy mitigated the expansion of effector cells in healthy mice (Figure 1E). Similarly, the combination of ImmTOR with an IL-2 mutein mitigated effector cell expansion following administration of high vector doses of AAV, which can cause hepatic inflammation^57^ (Suppl Figure 6B). One potential concern for Treg-selective IL-2 therapies is that in settings of inflammatory disease, activated effector T cells can transiently express IL-2Rα, leading to the formation of the high affinity IL-2Rαβγ^10, 11^. Indeed, following adoptive transfer of human PBMC into immunodeficient mice, a setting which can lead to GVHD, administration of the Treg-selective IL-2 fusion protein F5111 IC alone led to exaggerated expansion of effector T cells (Figure 2A). However, this expansion was prevented by the addition of ImmTOR. The increased expansion of effector cells observed in HuPBMC mice treated with F5111 IC alone correlated with exacerbation of disease in a HuPBMC GVHD model. Notably the addition of ImmTOR to F5111 IC significantly increased survival in this model (Figure 2B). Disease exacerbation has also been reported for the IL-2/JES6-1 anti-IL-2 antibody complex in a mouse model of inflammatory arthritis induced by infection with chikungunya virus^63^. Administration of IL-2/JES6-1 during active infection increased both Tregs and effector T cells resulting in disease worsening, while prophylactic administration in healthy mice prevented subsequent disease. Thus the activity of engineered IL-2 molecules may be dose-limited due to its effects on effector cells in settings of inflammation. The addition of ImmTOR increases the therapeutic window of Treg-selective IL-2 by restraining effector cell activation while synergistically increasing Tregs. Another potential concern related to engineered IL-2 molecules is their potential for immunogenicity. ImmTOR has been shown to inhibit immunogenicity of a variety of co-administered biologic therapies and could thus counteract potential anti-drug antibody responses^37^. Caution is warranted, as the combination of rapamycin with low dose IL-2 was reported to induce transient impairment of β cell function in a small clinical trial conducted in patients with Type 1 diabetes^64^. The authors speculated that the toxicity was related to IL-2, as β cell impairment was observed in patients that did not receive the full course of rapamycin and was most significant in the first month of therapy, concordant with IL-2 treatment. Our use of an engineered Treg-selective IL-2 molecules with long circulating half-life combined with ImmTOR could help mitigate potential toxicities associated with low dose cytokine administration.

The combination of ImmTOR with engineered Treg-selective IL-2 molecules showed potent synergistic activity in inhibiting antibody responses against an AAV gene therapy vector (Figure 5, Suppl Figure S5A). Currently, AAV gene therapies are limited to a single systemic administration due to development of high neutralizing antibody titers^57^. Even low titers of neutralizing antibodies may block transduction; thus, potent and durable inhibition of anti-capsid antibodies would be required to enable vector re-administration. ImmTOR+Fc.IL2m combination therapy also showed potent activity in preventing type 1 diabetes in NOD mice (Figure 6). In this model, ImmTOR+Fc.IL2m combined with a nanoencapsulated chromogranin A-insulin hybrid peptide provided the longest disease-free duration of activity, although ImmTOR and ImmTOR+Fc.IL2m administered in the absence of exogenous antigen also provided strong protection. Similarly, ImmTOR+Fc.IL2m showed significant activity in a mouse model of PBC, with marked reduction of peri-biliary mononuclear cell infiltrates, biliary hypercellularity and ductular ectasia (Figure 7). The addition of nanoencapsulated PDC-E2-ILD antigen further improved activity of ImmTOR+Fc.IL2m. Taken together, these results reinforce the importance of driving antigen-specific tolerogenic responses provided by co-administration of nanoencapsulated antigens. However, the efficacy of ImmTOR+Treg-selective IL-2 alone in models of both T1D and PBC suggests the possibility that this combination may be able to induce tolerogenic immune responses to endogenously expressed autoantigens in the context of autoimmune disease. In summary, our work demonstrates that combining ImmTOR with engineered Treg-selective IL-2 molecules provides a promising approach to mitigate pathogenic autoimmunity by leveraging both bystander suppression through expansion of total Tregs as well as inducing and expanding antigen-specific Tregs.

## Materials and Methods

### ImmTOR, other nanoparticles and IL-2 mutein molecules

Rapamycin containing nanoparticles (ImmTOR) were manufactured as described earlier^34, 35^. ImmTOR doses were based on rapamycin content ranging from 50 to 300 μg per mouse. Rapamycin (sirolimus) was manufactured by Concord Biotech (Ahmedabad, India). Antigen-containing nanoparticles (NP) were prepared using a water/oil/water (W/O/W) double-emulsion solvent evaporation method as described^35^. Briefly, PLGA (50:50) and pegylated polylactic acid (PLA-PEG) were dissolved in dichloromethane to form the oil phase. An aqueous solution of antigen (chicken ovalbumin or OVA protein, hybrid insulin peptide HIP6.9 (LQTLALNAARDP), or PDC-E2-ILD (amino acids 213-314)^65^ was then added to the oil phase and emulsified by sonication (Branson Digital Sonifier 250A). Following emulsification of the antigen solution into the oil phase, a double emulsion was created by adding an aqueous solution of polyvinyl alcohol and sonicating a second time. The double emulsion was added to a beaker containing PBS and stirred at RT for 4 h to allow the dichloromethane to evaporate. The resulting NPs were washed twice by centrifuging at 75,600 × g for 50 min at 4 °C followed by resuspension of the pellet in PBS. Concentration of extracted antigens was measured by HPLC. Dynamic Light Scattering (DLS) analysis of particle size and PDI was performed using a Malvern Zetasizer Nano-ZS ZEN 3600. All the nanoparticles loaded with antigens exhibited a particle size distribution ranging between 140-155 nm with a low polydispersity index (<0.15). Recombinant PDC-E2-ILD was manufactured by Genscript (Piscataway, NJ) using its proprietary E. coli expression system. Mouse IL2 mutein (Fc.IL2m) was constructed based on the sequence Fc.Mut24 published by Khoryati et al.^13^ The protein was manufactured by Genscript, using its proprietary CHO mammalian expression system. F5111 IC was produced as previously described^20^.

### Viruses

AAV8-SEAP was manufactured by SAB Tech (Philadelphia, PA, USA) using their proprietary helper plasmid and AAV8AAP plasmids, and the plasmid containing the gene of interest. The plasmids were transfected into adherent human embryonic kidney (HEK) 293 cells and harvested 72 hours after transfection by cell lysis. The clarified supernatant from the harvest was purified by CsCl_2_ gradient, and the AAV8-containing fraction was collected. The viral vector band was assayed by SDS–polyacrylamide gel electrophoresis (PAGE) gel and silver stain to determine a viral titer, which was then confirmed by qPCR using ITR-specific primers.

### Mice

Immunologically naïve, female C57BL/6 mice aged 36-52 days (or 17-18g) were purchased from Charles River Laboratories (Wilmington, MA). Similarly aged B6.Cg-Tg(TcraTcrb)425Cbn/J mice (also known as OT-II mice), expressing a T cell receptor (TCR) specific for chicken ovalbumin 323-339 peptide (OVA323-339) in the context of I-Ab resulting in OVA-specific CD4^+^ T cells were purchased from Jackson Laboratories (Bar Harbor, ME). Non-obese diabetic (NOD) NOD/ShiLtJ strain and FoxP3-IRES-mRFP (C57BL/6-Foxp3tm1Flv/J) mice co-expressing expressing the Foxp3 (forkhead box P3) gene with monomeric red fluorescent protein (mRFP) were also purchased from Jackson Laboratories. Human Tg-IL-2/IL-2Rα/IL-2Rβ mice carrying knock-out mutations for IL-2, and IL-2 receptor alpha and beta chains and expressing their human homologues were purchased from Biocytogen (Wakefield, MA). NCG mice (NOD-*Prkdc26^emCd52^Il2rg^em26Cd^*^22^/NjuCrl) carrying a mutation in *Sirpa* and knockouts of *Prkdc* and *Il2rg* genes and thus lacking functional/mature T, B, and NK cells, along with reduced macrophage and dendritic cell function were purchased from Charles River Laboratories. NCG mice were reconstituted with human PBMC from one of three different donors used in separate studies per manufacturer’s instructions. A similar strain, NSG (NOD.Cg-Prkdcscid Il2rgtm1Wjl/SzJ), carrying mutations in Prkdc and null allele of the IL2rg (IL2rgnull) lacking functional/mature T, B, and NK cells was purchased from Jackson Labs, and used in GVHD studies. NOD.c3c4 (NOD.B-(D3Mit93-D3Mit124)(D4Mit114-D4Mit142)/1112MrkJ) mice, known to spontaneously develop autoimmune liver disease and specifically, primary biliary cholangitis (PBC)62, 63 were purchased from Jackson Labs and then bred in-house. To minimize the potential effects of stress, mice were acclimated to the Animal Care Facility at Selecta for at least three days prior to injection. All the experiments were conducted in strict compliance with NIH Guide for the Care and Use of Laboratory Animals and other federal, state and local regulations and were approved by Selecta’s IACUC.

### Animal Injections

Mice were injected (i.v., tail vein or retro-orbital plexus) with ImmTOR nanoparticles in the effective range of 50-300 μg per mouse, or with NP-encapsulated protein or peptide antigens in the effective range of 0.05-5 µg per mouse, or with Fc.IL2m (i.p. or i.v., retro-orbital plexus) or F5111-IC (i.v.) in the effective range of 6.25-18.75 μg per mouse. Free ovalbumin (OVA) was administered i.v. in the effective range of 20-500 µg per mouse. AAV8-SEAP vector injected i.v. at specified doses. NOD (type 1 diabetes model) and NOD.c3c4 (PBC model) mice were treated with individual therapeutic components or their combinations as described in Figure Legends three or four times total at 28-day intervals.

### GVHD model

NSG mice (NOD.Cg-Prkdcscid Il2rgtm1Wjl/SzJ; Jackson Laboratory #005557) were irradiated with 1 Gy from an X-ray irradiator source and then reconstituted with 1×10^7^ human PBMC. The next day, mice were treated with a single dose of phosphate buffer saline (vehicle), ImmTOR (100 µg), F5111 IC (9 µg), or their combination. Animals were assessed for disease activity 3 times per week. Each animal was assessed for weight loss, posture, activity, fur texture, skin integrity, and paleness on a 2 grade scale as indicated below. Animals losing more than 20% weight or moribund were removed from the study.

| Criteria | Grade 0 | Grade 1 | Grade 2 |
| --- | --- | --- | --- |
| **Weight Loss** | <10% | from $\geq 10\%$ to $\leq 20\%$ | >20% |
| **Posture** | Normal | Hunching noted at rest | Severe hunching impairs movement |
| **Activity** | Normal | Mild to moderately decreased | Stationary unless stimulated |
| **Fur Texture** | Normal | Mild to moderate ruffling | Severe ruffling/poor grooming |
| **Skin Integrity** | Normal | Scaling of paws/tail | Obvious areas of denuded skin |
| **Paleness** | Normal | Slight paleness | Severe paleness |

### T1D model

NOD mice were enrolled in the study at 6-7 weeks of age with the first treatment at week 8. They were monitored weekly using standard glucometer strips, and mice showing glucose levels >250 mg/dL on 2/3 successive measurements were considered diabetic and those scoring >500 mg/dL twice or >600 mg/dL once were terminated. All animals in the study were terminated at 35 weeks.

### PBC model

NOD.c3c4 mice were enrolled in the study at 8 weeks and treated for the first time at 10 weeks of age. After termination at 24 weeks, livers were fixed, embedded in paraffin, sectioned, stained with hematoxylin and eosin and then slide images were taken. The resulting slides were evaluated by a certified veterinary pathologist, and microscopic findings were scored as follows: Grade 0 (normal): finding not present, Grade 1 (minimal): a focal, subtle, or trivial change, Grade 2 (mild): an easily identifiable change of limited severity and/or distribution, Grade 3 (moderate): an obvious change with normal tissue remaining, Grade 4 (marked): an extensive change that obliterates much of the normal tissue, Grade 5 (severe): a maximal change.

### Flow Cytometry (murine liver and spleen cell populations)

Immediately after euthanizing mice (at 1-14 days after initial treatment), livers and spleens were harvested and rendered into single cell suspensions. Livers were processed via collagenase 4 (Worthington, Lakewood, NJ) enzymatic digest according to manufacturer’s recommended protocol. Spleens were processed via mechanical passage through a 70 µm nylon mesh (ThermoFisher, Waltham, MA). Next, a red blood cell lysis step was performed for both liver and spleen suspensions for 5 min at room temperature in 150 mM NH_4_Cl, 10 mM KHCO_3_, 10 μM Na_2_-EDTA; washed in PBS, 2% bovine serum; then filtered again with a 70 µm nylon mesh. Cells were incubated 20 min on ice with anti-CD16/32 (Fc-block, clone 93, BioLegend, San Diego, CA) then stained with the following antibodies directed toward cell surface receptors: CD3e-BV421 (BioLegend, clone 145-2C11), CD4-PerCP-Cy5.5 (BioLegend, clone RM4-5), CD8a-BV510 (BioLegend, clone 53-6.7), CD25-PE-CF594 (BD, clone PC61), NK1.1-AF700 (BD, clone PK136), CD122-APC (BioLegend, clone TM-B1), and CD132-PE (BioLegend, clone TUGm2). Adoptively transferred human PBMC were stained with the following antibodies: CD4-PerCP-Cy5.5 (BioLegend, clone SK3), CD8a-APC-Cy7 (BioLegend, clone RPA-T8), CD56-BV421 BioLegend clone HCD56), CD3e-BV-510 (BioLegend, clone UCHT-1) and CD26-PE-CF594 (BD clone M-A251). After cell surface labeling cells were then fixed and permeabilized according to manufacturer’s recommended protocol using a FoxP3 Transcription Kit (eBioscience, San Diego, CA). The targeted intracellular markers were stained with FoxP3-PE (InVitrogen, Waltham, MA), clone FJK-16s, Ki67-A647 (BioLegend, clone 11F6) and Helios-PE-Cy7 (BioLegend, clone 22F6). Analysis was performed via FACSymphony A3 Cell Analyzer (BD Biosciences) with subsequent data analysis using FlowJo software (TreeStar, Ashland, OR).

### Whole animal imaging

FoxP3-IRES-mRFP mice were used to measure in vivo FoxP3 expression after treatment with Fc.IL2m alone or combined with ImmTOR. At various time-points post treatment, images were acquired of the left side dorsal aspect with the AMI Imaging System charge-coupled-device camera and analyzed with the Aura 4.0.7 software package (Spectral Instruments Imaging, Tucson, AZ). A region of interest (ROI) was manually selected based on signal intensity. The area of the ROI was kept constant, and the intensity was recorded as maximum [photons s-^1^ x cm^-2^ x sr^-1^ (steradian)] within a ROI.

### Methylation Analysis

Murine CD3^+^, CD4^+^CD3^+^ and CD4^+^CD25^+^ cells were isolated from splenocytes seven days post treatment via immunomagnetic bead selection (Miltenyi, Gaithersburg, MD) using either negative selection of untouched CD4^+^ T cells or positive selection of CD4^+^CD25^+^ T cells (both from Miltenyi). After careful supernatant removal, accurate cell counting (Countess, ThermoFisher), cell pellets were then snap frozen in liquid nitrogen, then stored on dry ice. Samples were then sent to EpigenDx (Hopkinton, MA) for subsequent targeted NextGen bisulfite sequencing panel analysis using their in-house FoxP3 methylation panel N4V1P15 analysis.

### Serum cytokines and FGF21 determination

Serum cytokine concentrations were determined using Meso Scale Discovery (MSD) U-PLEX 10-Assay SECTOR™ Plates, Linkers, and corresponding capture and detection antibody pairs. Plates were read using electrochemiluminescence detection on an MESO® QuickPlex SQ 120, with Discovery Workbench software (version 4.0.13) for analysis (MSD®, Gaitherburg, MD). Assays were performed according to manufacturer’s instructions, and without alterations to the recommended standard curve dilutions. Serum FGF21 concentration was determined by ELISA using the mouse/rat FGF21 commercial kit from R&D Systems (Minneapolis, MN). Serum samples were run at a 1:10 dilution.

### Concanavalin A Challenge Model

Concanavalin A (Con A) induced liver toxicity model was employed essentially as earlier described^36^. Briefly, mice were injected (i.v., r.o.) Con A at 12 mg/kg and then terminally bled at 6 or 12 hours post-challenge with cytokine levels in serum determined by MSD as described above and liver tissues collected simultaneously for single-cell suspension analysis by flow cytometry as described above or for hematoxylin-eosin staining followed by microscopic evaluation.

### Enzyme-linked immunosorbent assay for IgG against AAV8

The 96-well plates were coated overnight with AAV8, washed, and blocked on the following day, followed by sample incubation (1:40 diluted serum). Plates were then washed, and the presence of IgG was detected using anti-mouse IgG-specific horseradish peroxidase (HRP; 1:1500; Jackson ImmunoResearch, West Grove, PA, USA). The presence of rabbit anti-mouse IgG-specific HRP was visualized using trimethylboron substrate and measured using absorbance at 450 nm with a reference wavelength of 570 nm. The optical density (OD) observed is proportional to the amount of anti-AAV8 IgG antibody in a sample and was reported.

### Statistical Analysis

Statistical analyses were performed using GraphPad Prism 9.4.1. To compare the mouse experimental groups pairwise either multiple t test (for several time-points) or Mann-Whitney two-tailed test (for a single time-point; individual comparison of two groups presented within the same graph) were used. Significance is shown for each figure legend (* – p < 0.05, ** – p < 0.01; *** – p < 0.001; **** – p < 0.0001; not significant – p > 0.05). All data for individual experimental groups is presented as mean ± SD (error bars).

## Acknowledgements

We thank Drs. Daniel J. Campbell and Marc A. Gavin for sharing the sequence of Fc.IL2m.

## Author contributions

T.K.K. conceived the idea for the combination therapy, T.K.K. and P.O.I. designed the research and wrote the manuscript, M.F., A.M., G.R., and C.R. conducted the biological studies, T.C., M.F., G.R., and C.R, analyzed samples, N.N. and L.D. formulated the antigen nanoparticles, F.F. analyzed the nanoparticles, N.L. produced the protein products, D.VD and J.B.S. provided F5111 IC, P.G.T. analyzed the histology slides, T.K.K., P.O.I, S.L., T.C., M.F., G.R., and C.R, analyzed the data, T.K.K., P.O.I., D.VD, and J.B.S. edited the manuscript.

---

**Suppl. Fig. 1.**
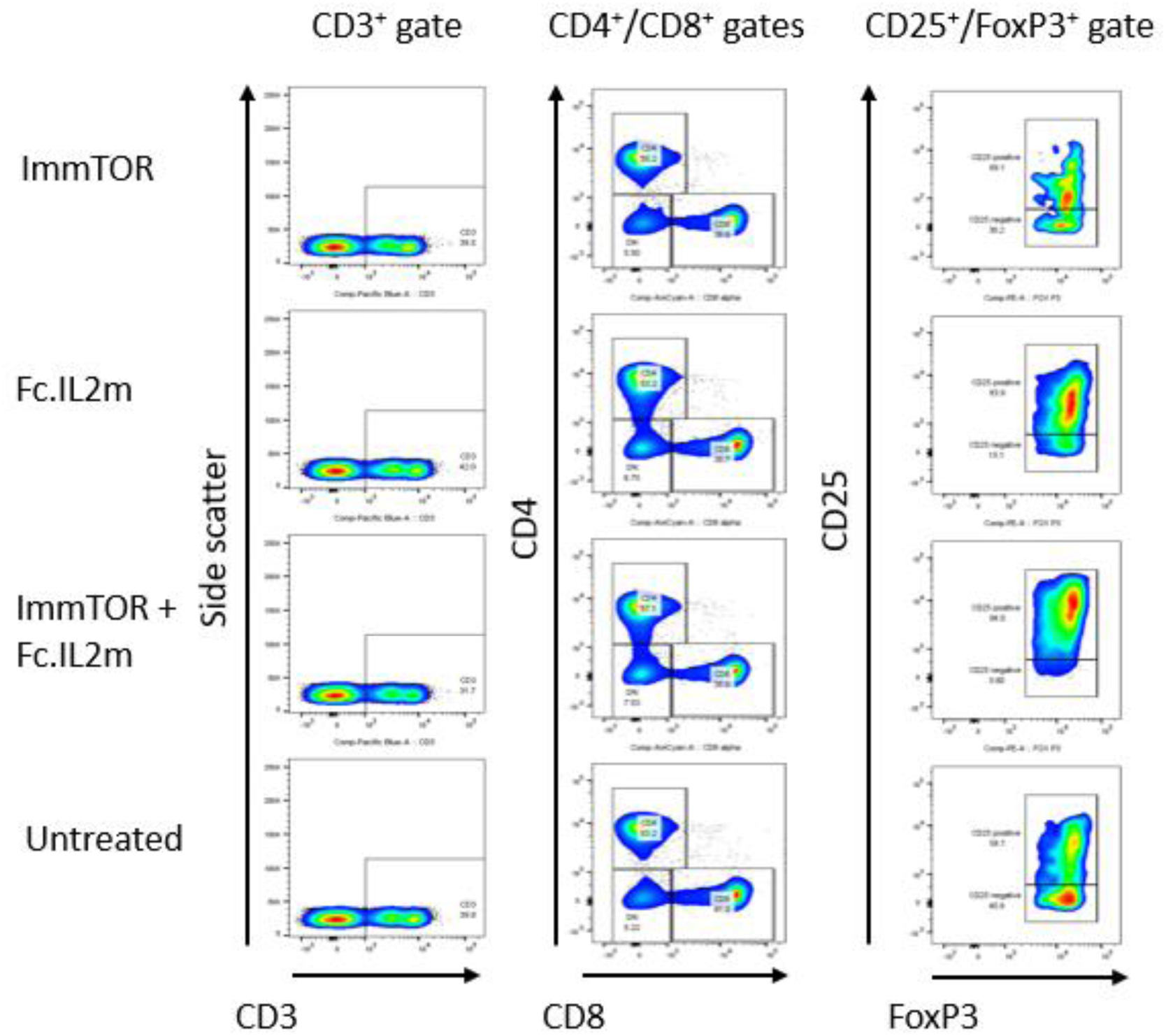
CD3, CD4, CD25 and FoxP3 gating strategies. Representative dot plots are shown for phenotyping of untreated mice and mice treated with ImmTOR, Fc.IL2m (IL-2 mut), or ImmTOR+Fc.IL2m combination therapy.

**Suppl. Fig. 2.**
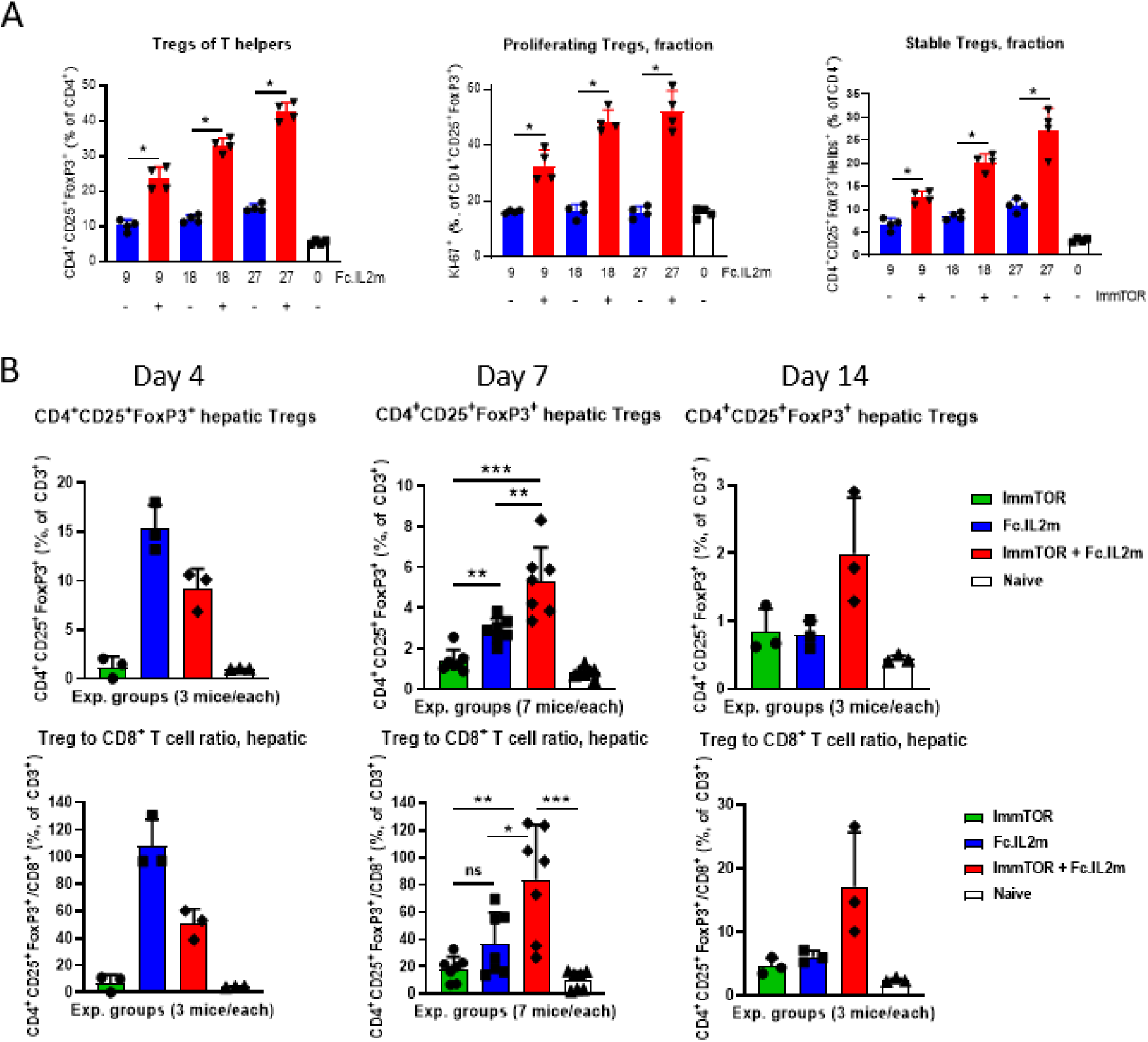
Treg induction by Fc.IL2m +/- ImmTOR in spleen and liver. **A.** Dose-dependence of Treg induction by Fc.IL2m alone or combined with ImmTOR. Groups of mice (n=4 per cohort) were treated and splenic Tregs were analyzed as described in Figure 1C. Fractions of Tregs out of total T helpers and of proliferating (Ki67^+^) and Helios^+^ stable Tregs out of total Tregs are shown. Error bars indicate mean +/- SD. **B.** Dynamics of hepatic Treg induction by combination treatment with ImmTOR and IL-2 mutein. Groups of mice (n=3-7 per cohort per time-point) were treated with ImmTOR, Fc.IL2m or the combination thereof. Livers were harvested on Day 4, 7, or 14, as indicated, processed to single-cell suspension, stained, and analyzed by flow cytometry. Graphs represent summaries of 4 independent experiments. Total Treg numbers and ratio of Treg-to-CD8^+^ T cells are shown. Error bars indicate mean +/- SD. Statistical significance: * p < 0.05, ** p < 0.01, *** p < 0.001.

**Suppl. Fig. 3.**
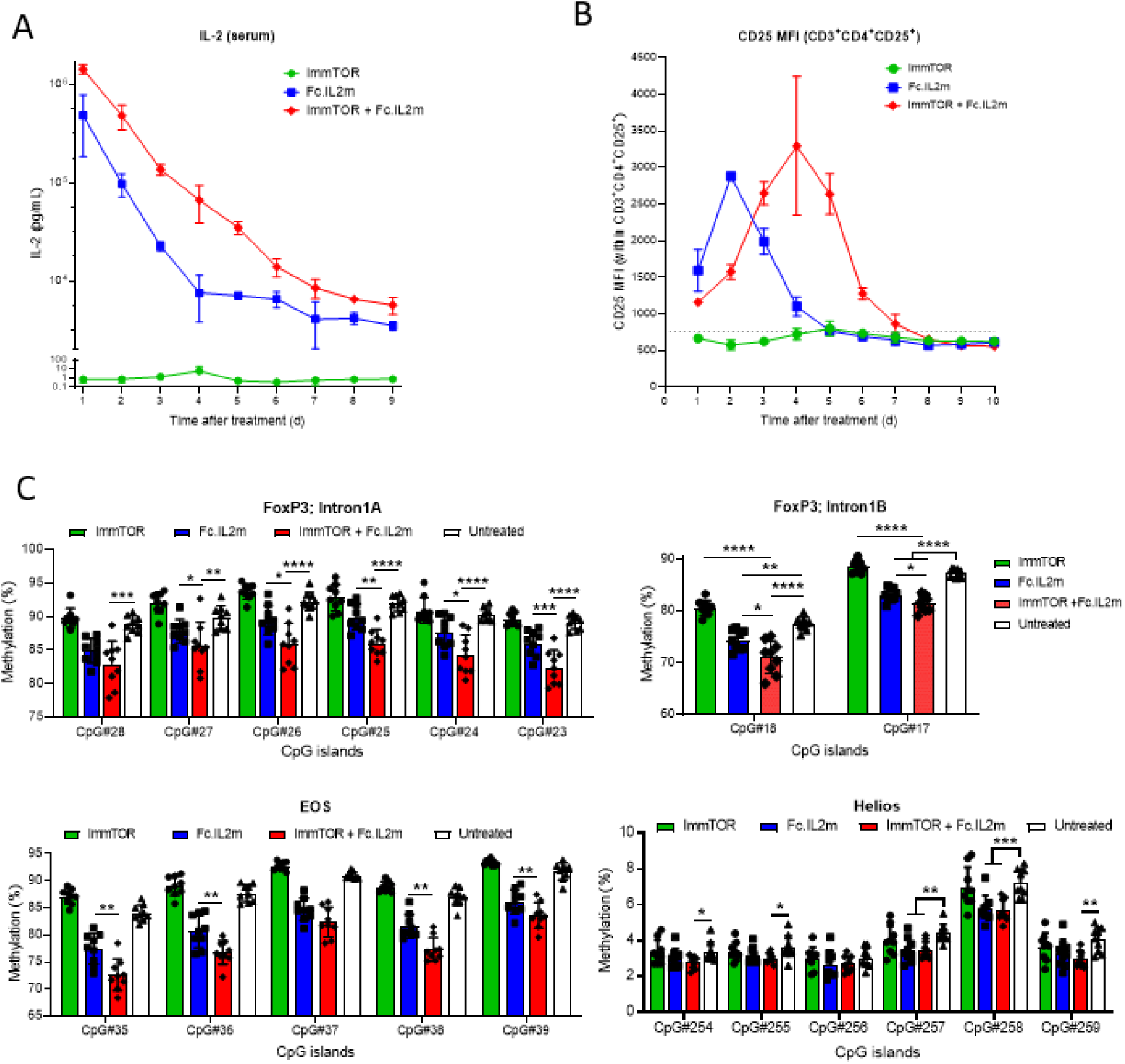
Treg responses to Fc.IL2m +/- ImmTOR. **A**, **B**. Dynamics of circulating Fc.IL2m **(A)** and CD4^+^ T cell IL-2Rα (CD25) expression (**B**) after treatment with ImmTOR, Fc.IL2m, or the combination thereof. Groups of mice (n=3-6 per group for each timepoint) were treated with Fc.IL2m and/or ImmTOR at the indicated doses. The graphs represent summaries of 3 independent experiments. Error bars indicate mean +/- SD. **C**. Demethylation of Treg-specific genes after treatment with ImmTOR, Fc.IL2m or their combination. Groups of mice (n=9 per group) were treated with Fc.IL2m and/or ImmTOR at the doses indicated, CD4^+^ T cells were isolated after 7 days, and status of methylation within FoxP3, EOS, and Helios genes was assessed within multiple CpG islands as indicated. The graphs represent summaries of 2 independent experiments. Statistical significance: * p < 0.05, ** p < 0.01, *** p < 0.001, **** p < 0.0001.

**Suppl. Fig. 4.**
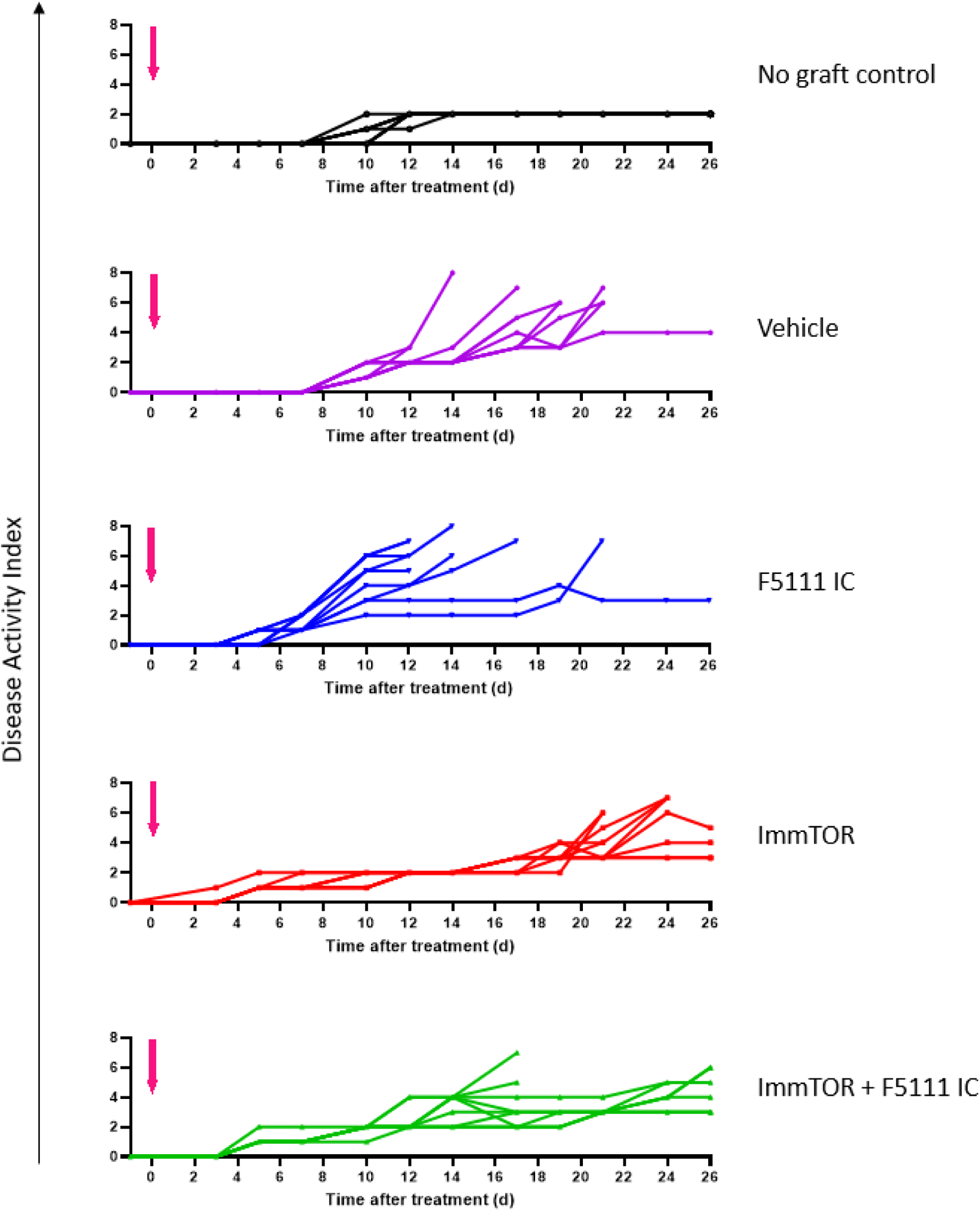
ImmTOR improves GVHD disease scores. NSG mice were irradiated, reconstituted with HuPBMC and treated as described in Figure 2B legend. Disease activity index (DAI) was assessed three times per week, as described in Materials and Methods.

**Suppl. Fig. 5.**
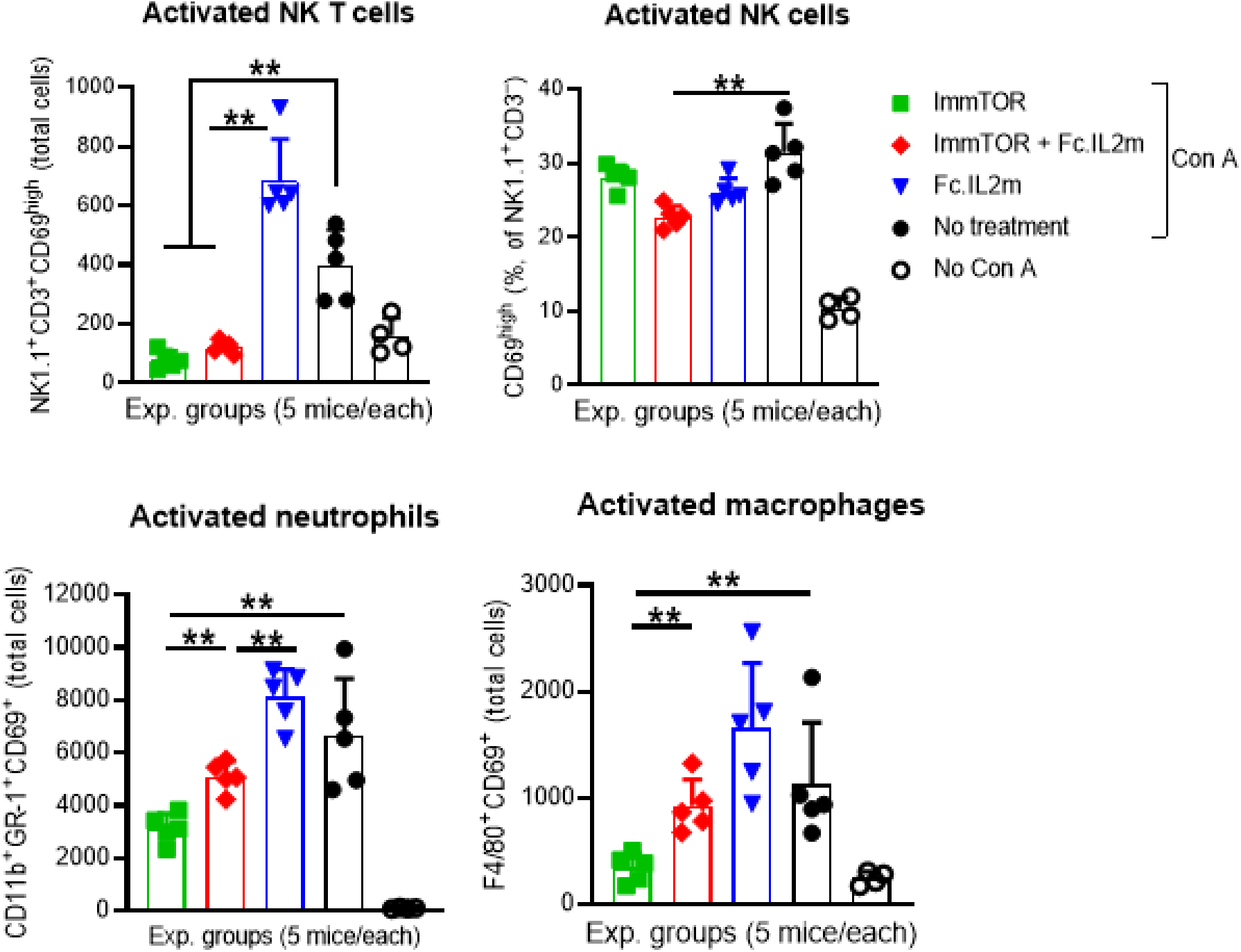
Effect of ImmTOR and IL-2 mutein on activation of hepatic NK, NKT, neutrophils, and macrophages after treatment with concanavalin. Female C57BL/6 mice (n=5 per cohort) were either left untreated or treated with ImmTOR (200 µg) and Fc.IL2m (9 µg) individually or in combination. Mice were challenged 4 days later with 12 mg/kg of concanavalin A (Con A), as described in Figure 3. At 12 hours after Con A challenge, serum was drawn for cytokine quantification and livers were harvested and hepatic T cells were analyzed by flow cytometry. (**A**) Fractions or total cell numbers of activated (CD69^+^ or CD69^high^) NK (CD3^−^NK1.1^+^), NKT (CD3^+^NK1.1^+^) cells, neutrophils (CD11b^+^GR-1^+^), and macrophages (F4/80^+^) are shown. (**B**) Serum concentrations of IFN-γ, IL-6 and CXCL1. (**C**) Serum concentrations of FGF21. Error bars indicate mean +/- SD. Statistical significance: ** p < 0.01.

**Suppl. Figure 6.**
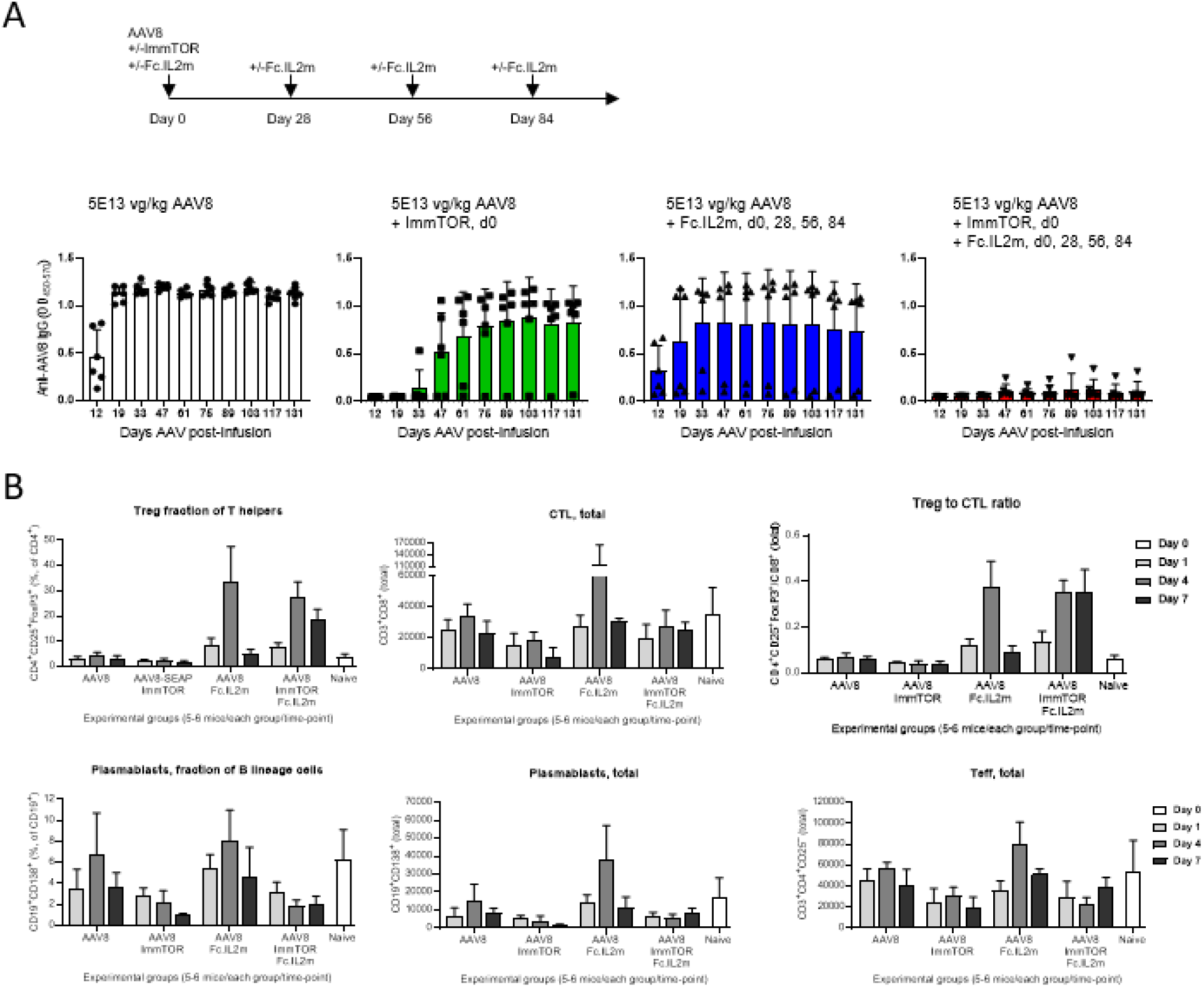
Mitigation of antibody response to high AAV vector dose by combination treatment with ImmTOR and IL-2 mutein. (**A**) C57BL/6 mice (n=6 per cohort) were treated with a single high vector dose of 5E13 vg/kg on Day 0 with or without ImmTOR (200 µg) and/or Fc.IL2m (9 µg). Groups treated with Fc.IL2m received additional Fc.IL2m doses on Days 28, 56, and 84. Anti-AAV8 IgG antibodies were assessed via serum blood draw on various days as indicated. (**B**) Spleens were harvested at the timepoints shown and processed into single-cell suspensions that were analyzed by flow cytometry. Populations are presented as fractions, absolute cell numbers, and relative ratios. Graphs are the summary of 2 identical studies with similar results. Error bars indicate mean +/- SD. Statistical analyses shown in Supplementary Table ST2.

**Supplementary Table ST1.**
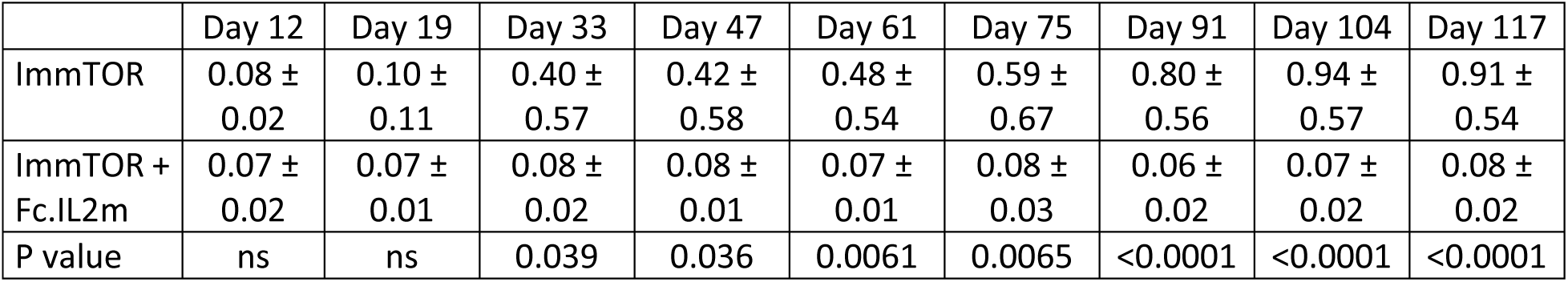
Comparison of anti-AAV IgG levels in mice treated with ImmTOR alone (50-200 µg) vs. animals treated with ImmTOR (50-200 µg) combined with Fc.IL2m (9 µg) as described in legend to Figure 5. Mean values ± SD are shown for all time-points and statistical significance indicated (unpaired t-test; ns – not significant).

**Supplementary Table ST2.** Statistical significance of differences between all the splenocyte cell populations (fractions, absolute numbers and their ratios) shown in Supplementary Fig. 6B. The values for the group of mice treated with AAV and ImmTOR and Fc.IL2m combination vs. all other experimental groups at all time-points is shown. * p < 0.05, ** p < 0.01, ns – not significant.

|  | AAV |  |  | AAV + ImmTOR |  |  | AAV + IL-2mut |  |  |
| --- | --- | --- | --- | --- | --- | --- | --- | --- | --- |
|  | Day 1 | Day 4 | Day 7 | Day 1 | Day 4 | Day 7 | Day 1 | Day 4 | Day 7 |
| Treg fraction (% of CD4<sup>+</sup>) | ** | ** | ** | ** | ** | ** | ns | ns | ** |
| CTL, total | ns | ns | ns | ns | ns | ** | ns | ** | * |
| Treg:CTL ratio | ** | ** | ** | ** | ** | ** | ns | ns | ** |
| Plasmablasts (% of CD19<sup>+</sup>) | ns | ** | * | ns | ns | ** | * | ** | * |
| Plasmablasts, total | ns | * | ns | ns | ns | ** | * | ** | ns |
| T effectors, total | * | ** | ns | ns | ns | * | ns | ** | * |

